# Gba1 deletion causes immune hyperactivation and microbial dysbiosis through autophagic defects

**DOI:** 10.1101/2022.12.15.520449

**Authors:** Magda Luciana Atilano, Alexander Hull, Catalina-Andreea Romila, Mirjam L Adams, Jacob Wildfire, Enric Ureña, Miranda Dyson, Jorge Ivan-Castillo-Quan, Linda Partridge, Kerri J. Kinghorn

**Author notes:** These authors contributed equally to the work (Magda Luciana Atilano, Alexander Hull).

## Abstract

Mutations in the *GBA1* gene cause the lysosomal storage disorder Gaucher disease (GD) and are the greatest genetic risk factor for Parkinson’s disease (PD). Communication between gut and brain and immune dysregulation are increasingly being implicated in neurodegenerative disorders such as PD. Here, we show that flies lacking the *Gba1b* gene, the main fly orthologue of *GBA1*, display widespread innate immune up-regulation, including gut inflammation and brain glial activation. We also demonstrate gut dysfunction in flies lacking *Gba1b*, with increased intestinal transit time, gut barrier permeability and microbiome dysbiosis. Remarkably, modulating the microbiome of *Gba1b* knockout flies, by raising them under germ-free conditions, can partially ameliorate lifespan, locomotor and some neuropathological phenotypes. Lastly, direct stimulation of autophagy by rapamycin treatment achieves similar beneficial effects. Overall, our data reveal that the gut microbiome drives systemic immune activation in *Gba1b* knockout flies and that reducing innate immune response activation either by eliminating the microbiota or clearance of immunogens by autophagy may represent potential therapeutic avenues for *GBA1-*associated neurodegenerative disease.

## Introduction

The *GBA1* gene encodes the lysosomal enzyme glucocerebrosidase (GCase), responsible for the hydrolysis of the key membrane sphingolipid glucosylceramide (GluCer) to produce glucose and ceramide. Bi-allelic mutations in the *GBA1* gene are known to cause Gaucher disease (GD), whereas heterozygous mutations represent a major genetic risk factor for Parkinson’s disease (PD), conferring an approximately 20-fold increased risk (Migdalska-Richards and Schapira, 2016, Schapira, 2015). GD is a multi-systemic disorder characterised by the lysosomal accumulation of GluCer within macrophages of the liver, spleen, bone marrow, and other tissues, causing widespread organ dysfunction and inflammation (Cox, 2010). To date there are no effective disease-modifying therapies targeting the neuropathology in GD (Mistry et al., 2017). As current GD therapies fail to permeate the blood-brain barrier, targeting a peripheral process to revert of prevent neurodegenerative pathologies may be beneficial. PD is characterised neuropathologically by the presence of intraneuronal inclusions called Lewy bodies, composed predominantly of aggregated α-synuclein (αSyn), which forms higher-order aggregates in the presence of GluCer, leading to further lysosomal dysfunction (Zunke et al., 2018). Another pathological hallmark is dopaminergic neuronal loss in the substantia nigra, where widespread GCase deficiency is also observed (Alcalay et al., 2015, Gegg et al., 2012, Spillantini et al., 1998).

There is growing evidence to suggest that the gut-brain axis is involved in the development of PD, and that the gut may act as an initiating site for PD pathology (Elfil et al., 2020, Parashar and Udayabanu, 2017). Moreover, studies spanning clinical and basic science research have demonstrated both central and peripheral immune changes in PD and GD, implicating the immune system in gut-brain axis communication. Multiple genetic studies have linked PD risk to immune-related pathways and pathogenic variants in immune genes (e.g. Toll-like receptor (TLR-4), Tumour Necrosis Factor (TNF)-α and Human Leukocyte Antigen-DR (HLA-DR) (Chu et al., 2012, Holmans et al., 2013, International Parkinson Disease Genomics et al., 2011, Zhao et al., 2015) Furthermore, impaired autophagy in macrophages derived from peripheral monocytes of GD patients leads to increased secretion of the pro-inflammatory cytokines interleukin (IL)-6 and IL-β in association with inflammasome activation (Aflaki et al., 2016). Despite the growing evidence linking immune dysfunction to sporadic PD and GD, the precise role of immune responses in gut-brain cross-talk in *GBA1*-PD and neuronopathic GD is yet to be elucidated.

In *Drosophila*, the innate immune response is initiated by pattern recognition receptors (PRRs), which recognize pathogen associated molecular patterns (PAMPs) on the surface of microbes to induce the conserved immune signal transduction pathways IMD, Toll and JAK/STAT (Buchon et al., 2014, Lemaitre and Hoffmann, 2007). Activation of innate immune pathways results in the up regulation of a cadre of structurally diverse antimicrobial peptides (AMPs) with bactericidal properties. Nevertheless, chronic immune pathway activation is deleterious to health and thus activation of the innate immune pathways must be tightly regulated (Cao et al., 2013). Autophagy is a biological process responsible for the removal of damaged organelles, aggregated protein, lipids and PAMPs. During autophagy the cellular material to be degraded is sequestered into double-membraned vesicles called autophagosomes. In a later stage, these structures fuse with lysosomes containing lysosomal hydrolases which degrade the sequestered cargo.

To study immune and gut pathologies in *GBA1*-associated disease, we utilised a *GBA1* (*Gba1b*) knockout Drosophila model, which has been described to recapitulate GD and PD pathology (Kinghorn et al., 2016). This model displays decreased lifespan and age-dependent locomotor defects, underpinned by elevated levels of the substrate GluCer, and an increase in the size and number of lysosomes. These abnormalities are associated with autophagy impairment and neurodegeneration. Here, we describe widespread innate immune activation and gut dysfunction in this *Gba1b* knockout fly model. Remarkably, we report that the intestinal microbiota is necessary for immune activation and that eradication of the altered gut microbiome in *Gba1b* knockout flies improves lifespan, locomotor and gut phenotypes. Moreover we demonstrate that direct stimulation of autophagy with rapamycin is sufficient to reduce immune activation and partially rescues GCase-deficient lifespan phenotype. Together, these results suggest that gut microbes play an important role in triggering the immune response in GD and *GBA1*-PD, and that targeting the gut microbiome or improving the clearance of PAMPs may represent a potential therapeutic strategy in these diseases.

## Results

### Loss of *Gba1b* results in innate immune activation

We have previously developed a *Drosophila* model of GCase deficiency by knocking out the two fly *Gba1* genes, *Gba1a* and *Gba1b* (Figure 1A), together or separately using homologous recombination (Kinghorn et al., 2016). Loss of *Gba1b* results in locomotor defects, reduced lifespan and neurodegeneration (Kinghorn et al., 2016). *Gba1b* display widespread expression and is thus the predominant *GBA1* fly orthologue. It is significantly expressed in the nervous system and heart, with strongest expression levels in the fat body (FlyAtlas2) (Leader et al., 2018). Several studies have identified that loss of *Gba1b*, but not *Gba1a*, recapitulates PD and GD phenotypes (Cabasso et al., 2019, Davis et al., 2016, Kinghorn et al., 2016, Maor et al., 2016). Our study therefore focused on flies lacking the *Gba1b* gene (*Gba1b*^*-/-*^).

**Figure 1.**
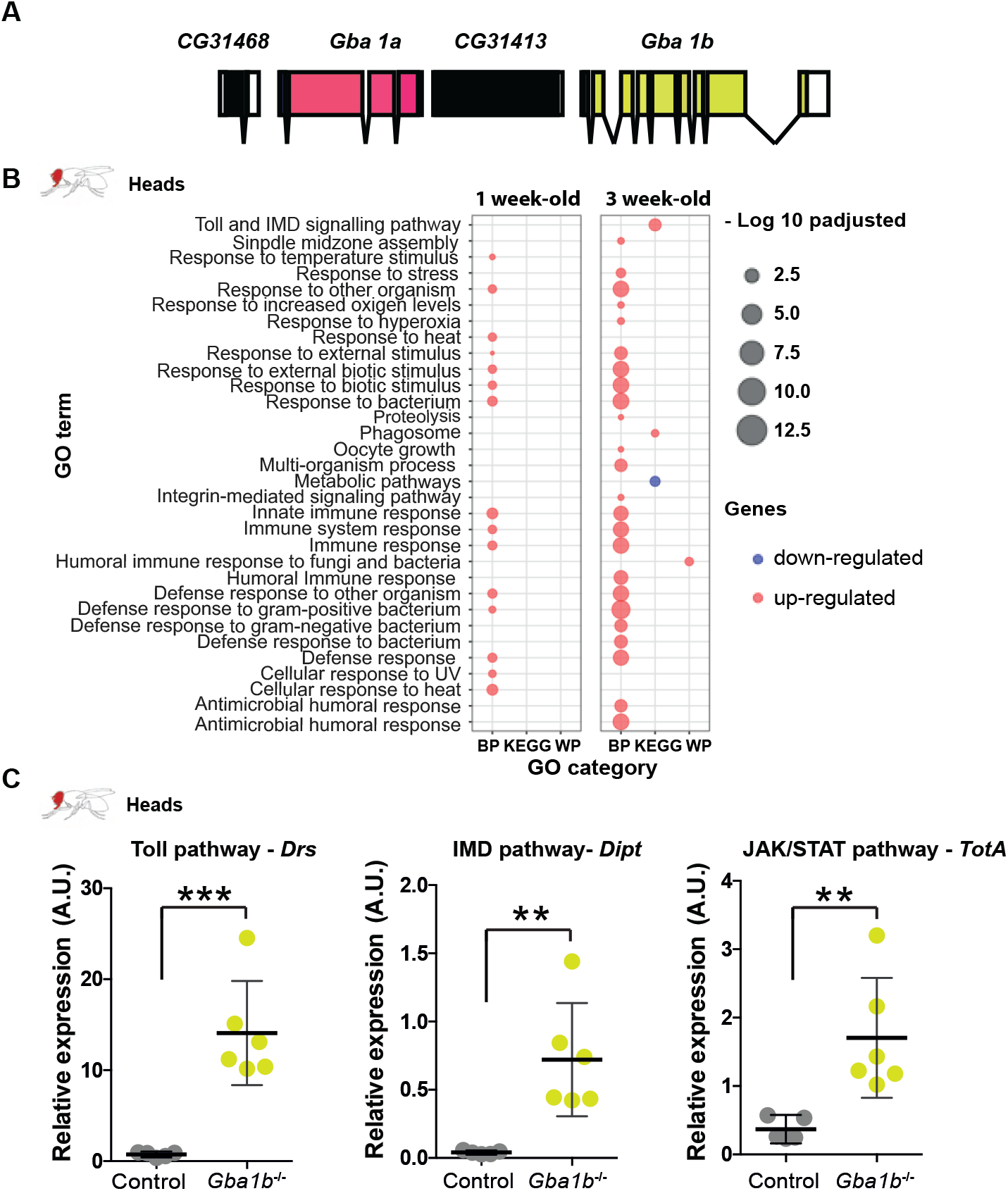
GCase deficiency results in up-regulation of innate immune pathways in the fly head. **(A)** Schematic representation of the two *Drosophila Gba1* gene loci, *Gba1a* and *Gba1b*. **(B)** Functional enrichment of the up-regulated and down-regulated genes in the heads of 1- and 3-week-old *Gba1b-/-* flies relative to controls. Visualization of all the significant GO-terms (adjusted p-values <0.05) for Biological Process (BP), KEGG pathway (KEGG) and Wiki pathway (WP) GO-categories demonstrates strong up-regulation of innate immune pathways. The size of the dots represents -log10 p-adjusted values for the GO-term enrichments. **(C)** Quantitative RT-PCR confirms up-regulation of the Toll (*Drs)* (***p=0.0004), IMD (*Dipt)* (**p=0.0041) and JAK-STAT (*TotA*) (**p=0.0069) reporter genes in 3-week-old *Gba1b*^*-/-*^ fly heads compared to controls. All target gene expression levels are normalized to tubulin. P-values were derived using an unpaired t-test and are presented as mean ± 95% confidence intervals, n=5-6 per genotype.

In order to identify molecular and biological processes contributing to the observed phenotypes in *Gba1b*^*-/-*^ flies, we performed RNA-sequencing analysis on the heads of 1- and 3-week-old *Gba1b*^*-/-*^ and age-matched controls. This revealed that 207 genes were differentially expressed in *Gba1b*^*-/-*^ fly heads compared to controls at both time points, with more genes differentially expressed at 3 weeks compared to 1 week (S1). Gene ontology analysis of both up- and down-regulated genes at both time points revealed significant gene enrichment for up-regulated genes mapping to pathways and biological processes related to the innate immune system, with greater up-regulation at 3 weeks (Figure 1B and S1). Many of these significantly expressed genes are key components of the Toll, IMD and JAK-STAT innate immune pathways (Figure S1).

Quantitative RT-PCR (qRT-PCR) was used to independently examine innate immune gene expression. The antimicrobial peptide (AMP) *Drosomycin* (*Drs*), a reporter gene of the Toll pathway, was significantly up-regulated in the heads of 3-week-old *Gba1b*^*-/-*^ flies compared to controls (Figure 1C). Similar changes were seen in the expression levels of *Diptericin* (*Dipt*), an AMP downstream of the IMD pathway and *TotA*, a reporter gene of the JAK-STAT pathway (Figure 1C). Interestingly, a similar increase in *Dipt* and *TotA* gene expression, but not in *Drs* levels, was seen in the headless bodies of aged *Gba1b*^*-/-*^ flies, confirming peripheral innate immune activation (S2).

### GCase deficiency leads to brain glial activation and immune up-regulation in the fat body and gut

Following on from the observation that local innate immune pathways are activated in the heads and bodies of *Gba1b*^*-/-*^ flies, the spatial pattern of the immune responses was studied in more detail. Given the presence of immune-responsive fat body tissue within the head, evidence of neuroinflammation was probed in the *Gba1b*^*-/-*^ fly brain by examining the highly conserved glial immune pathway involving Draper. Draper is a phagocytic recognition receptor on the surface of engulfing glia in flies and is required for the glial phagocytic removal of axonal debris following axonal injury (Doherty et al., 2009, MacDonald et al., 2006). Consistent with the widespread innate immune activation, increased *Draper* gene expression was observed in the heads of 3-week-old *Gba1b*^*-/-*^ flies (Figure 2A). Draper protein levels were also increased in the *Gba1b*^*-/-*^ fly brain on immunostaining (Figure 2B) and on Western blot analysis of fly heads (Figure 2C).

**Figure 2.**
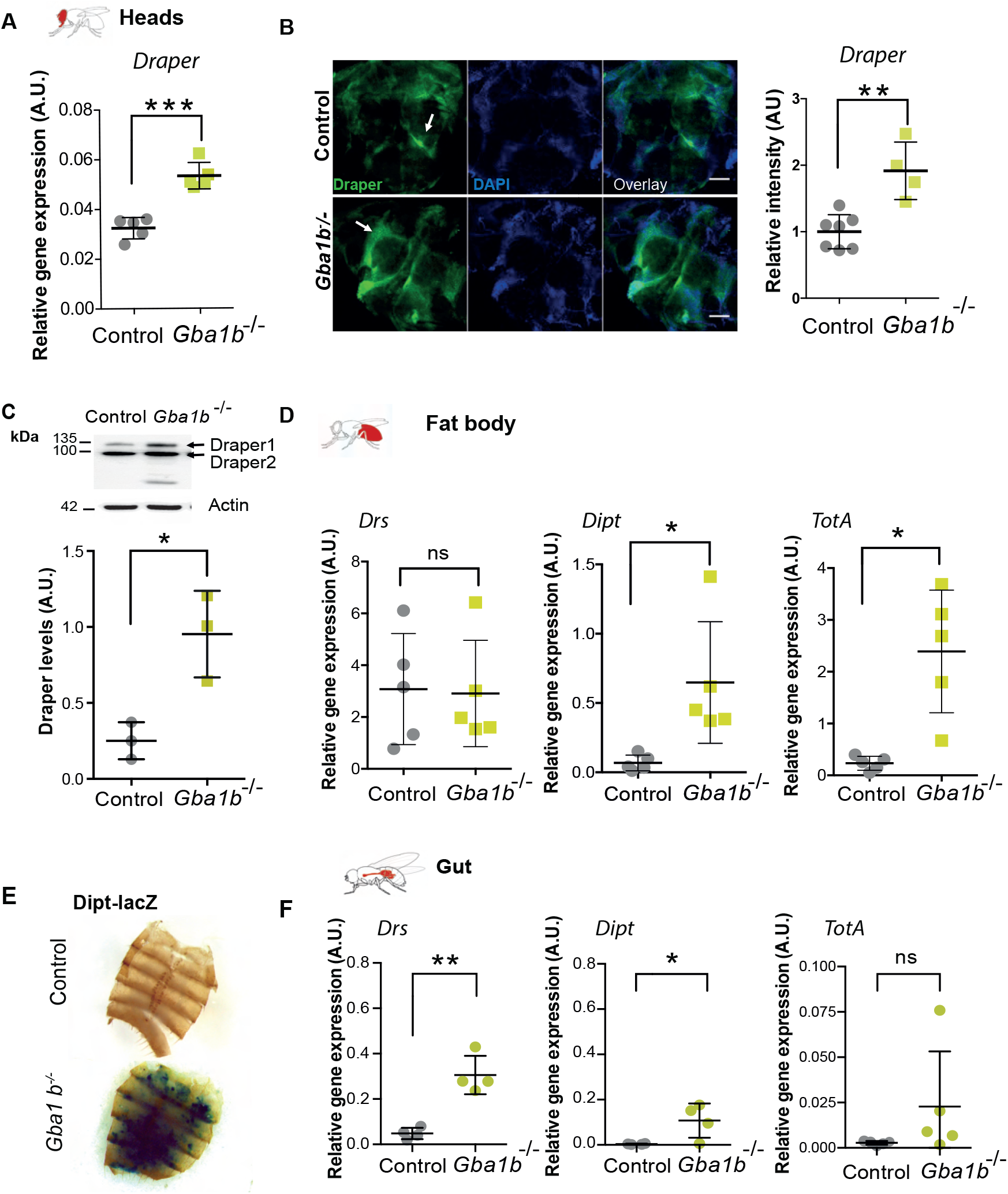
*Gba1b*^*-/-*^ flies display glial activation in the brain and immune activation in the fat body and gut. **(A)** Quantitative RT-PCR analysis demonstrates increased *Draper* gene expression in the heads of 3-week-old *Gba1b*^-/-^ flies compared to controls (***p=0.0001). Significance was assessed by unpaired t-test and presented as mean ± SD, n=5 per genotype. **(B)** Draper immunofluorescence (green channel) is increased in 3-week-old *Gba1b*^-/-^ fly brains compared to age-matched controls (**p=0.0015). White arrows show Draper localization in the antennal lobes. Scale bars, 50 µm. Representative images are shown. Significance was assessed by unpaired t-test and presented as mean ± SD, n=4-7 per genotype. **(C)** Western blot analysis confirms increased Draper I protein levels in the heads of *Gba1b*^-/-^ flies compared to controls. Draper levels are shown normalised to actin. Significance was assessed by unpaired t-test (*p=0.0354) and presented as mean ± SD, n=3 biological repeats per genotype. **(D)** Quantitative RT-PCR analysis confirms up-regulation of the IMD (*Dipt)* (*p=0.041) and JAK-STAT (*TotA*) (*p=0.0147) reporter genes in the pooled dissected fat bodies of 3-week-old *Gba1b*^*-/-*^ flies compared to controls. All target gene expression levels are normalized to tubulin. P-values were derived using an unpaired t-test and are presented as mean ± SD, n=5 per genotype. **(E)** Expression of the IMD reporter, Dipt-LacZ, in CRIMIC inserted *Gba1b* null flies revealed strong LacZ staining in the dissected fat body tissue. Representative images are shown. **(F)** The expression of the Toll innate immune pathway reporter gene, *Drs*, is increased in the midgut of 3-week-old *Gba1b*^*-/-*^ flies compared to controls (**p=0.0011). The IMD (*Dipt)* reporter gene is also significantly increased (*p=0.0334), while JAK-STAT reporter *TotA* is not significantly elevated (p=0.1801). All target gene expression levels are normalized to tubulin. P-values were derived using unpaired t-test and are presented as mean ± SD, n=4-5 per genotype.

Innate immune responses were then assessed in the fat body, the main site of AMP production. In keeping with IMD and JAK-STAT pathway up-regulation in the bodies of *Gba1b*^*-/-*^ flies, there was a significant increase in *Dipt* and *TotA* gene expression in dissected fat body tissue (Figure 2D). This was corroborated by the co-expression of an IMD reporter construct, Dipt-LacZ. This revealed strong diffuse LacZ staining in the fat body tissue of flies lacking *Gba1b*, consistent with IMD pathway activation (Figure 2E). An increase in innate immune signalling in the intestinal tissue was also observed. *Drs* and *Dipt* gene expression were significantly elevated in the midgut tissue of *Gba1b*^*-/-*^ flies compared to controls (Figure 2F). In addition, there was a trend towards up-regulation of *TotA* expression, although this did not reach statistical significance (Figure 2F).

It has recently been demonstrated that *Gba1b* is expressed predominantly within glia, rather than neurons, in the fly brain (Wang et al., 2022). Using the same CRIMIC *Gba1b* line, in which *Gba1b* gene expression is disrupted, we co-expressed UAS-mCherry.nls under the *Gba1b* endogenous promoter (Lee et al., 2018). mCherry expression almost exclusively overlapped with the glial marker repo, with little overlap with the neuronal marker elav, confirming the glial-specific expression of *Gba1b* (S3A). Interestingly, we also observed strong mCherry expression within the fat body and gut tissues (S3B and S3C). This suggests that the increased innate immune response observed could arise directly from *Gba1b* dysfunction within the tissues where the gene is expressed.

### *Gba1b* deficient flies exhibit increased intestinal transit time and increased gut wall permeability

Given the innate immune activation in the gut of *Gba1b*^*-/-*^ flies, intestinal pathophysiology was explored. PD patients exhibit intestinal inflammation, and gastrointestinal (GI) abnormalities such as constipation are a common symptom, often preceding motor defects by several years in PD (Devos et al., 2013). Changes in intestinal physiology were identified in *Gba1b*^*-/-*^ flies by analysing the rate of faeces production following the feeding of food supplemented with 0.5% bromophenol blue (BPB). Aged *Gba1b*^*-/-*^ flies exhibited a delay in intestinal transit time, with less blue faecal deposits produced in the first few hours after consuming BPB food compared to controls (Figure 3A). We also noted that the faecal content of control flies was more concentrated, with a high proportion of oblong shaped ROD deposits compared to non-ROD circular faecal deposits, whilst *Gba1b*^*-/-*^ flies almost exclusively produced non-ROD deposits (Figure 3A). The latter represent faeces with a higher water content (Cognigni et al., 2011). Taken together, our results show delayed intestinal emptying and altered faecal content in *Gba1b*^*-/-*^ flies, in keeping with the gastrointestinal dysfunction seen in PD. The altered intestinal transit was not due to reduced feeding, as a CAFE assay revealed no significant difference in blue food consumption between aged *Gba1b*^*-/-*^ flies and age-matched control flies (Figure S4).

**Figure 3.**
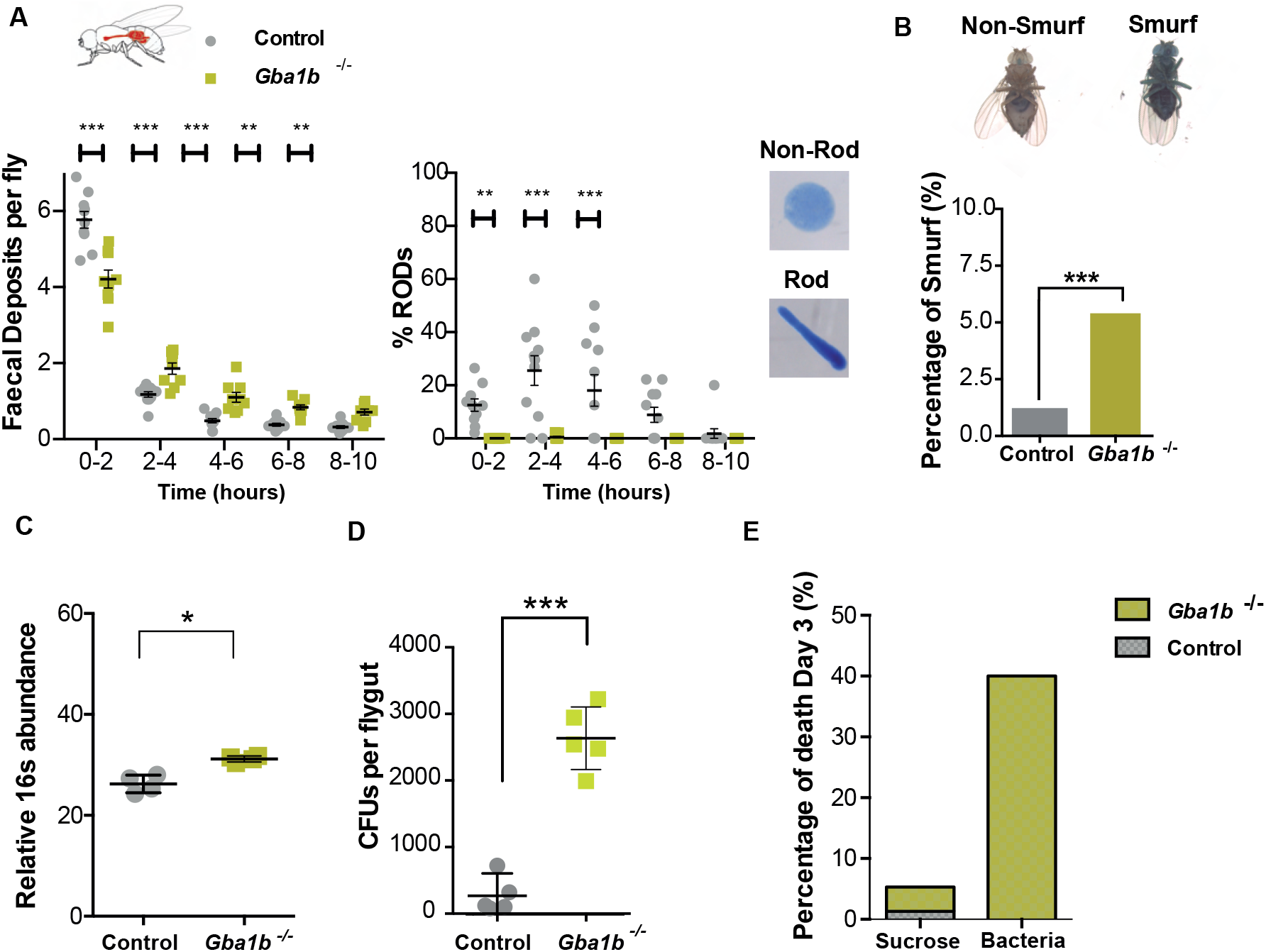
*Gba1b*^*-/-*^ flies show reduced intestinal transit and an altered gut microbiome. **(A)** The rate of intestinal transit is decreased in 3-week-old *Gba1b*^*-/-*^ flies compared to controls as assessed by the number of faecal deposits over time. *Gba1b*^*-/-*^ flies produce almost exclusively non-ROD faecal deposits. **(B)** Assessment of gut permeability using a Smurf assay revealed that there is an increase in the number of Smurf flies among aged *Gba1b*^-/-^ flies compared to controls, suggesting increased gut wall permeability (*Gba1b*^-/-^ vs control, ***p=0.0002;). P-values were calculated using X^2^ (chi-squared) tests with Yates’ correction. **(C)** Quantitative RT-PCR-based 16S rRNA gene abundance was significantly higher in the guts of *Gba1b*^*-/-*^ than in control flies. Significance was assessed by Mann-Whitney test (*p=0.0286), data presented as mean ± SD, n=4 per genotype. **(D)** Colony forming units (CFUs) were significantly increased in the guts of 3-week-old *Gba1b*^*-/-*^ flies compared to age-matched controls (p<0.001; unpaired t-test, result presented as mean ± SD, n=5 per genotype). **(E)** Oral infection with *Lactobacillus plantarum* resulted in increased percentage of mortality among *Gba1b*^*-/-*^ flies but not in control flies.

In addition to GI dysfunction, compromised intestinal barrier integrity has been reported in PD patients (Forsyth et al., 2011) (Schwiertz et al., 2018). We therefore employed a Smurf assay to assess gut permeability in *Gba1b*^*-/-*^ flies. This involved feeding with a non-absorbable blue food dye (Regan et al., 2016), and measuring the proportion of flies that display the presence of the blue dye outside of the gut, resulting in a blue Smurf appearance. Smurf flies were more abundant in the aged *Gba1b*^*-/-*^ population relative to age-matched controls, demonstrating compromised gut barrier integrity (Figure 3B).

### Flies lacking *Gba1b* exhibit an altered intestinal microbiome

Changes in the gut microbiome have been described in PD patients as a precursor to neuropathology (Dutta et al., 2019, Heller, 2018, Lin et al., 2019). In view of the intestinal immune activation and gut dysfunction in *Gba1b*^-/-^ flies, we performed quantitative RT-PCR analysis using 16S universal primers and bacterial culture on midgut tissue from aged *Gba1b*^*-/-*^ flies and controls to assess changes in microbiota load. A significant increase in overall bacterial load by quantitative RT-PCR was observed in midguts from *Gba1b*^*-/-*^ flies compared to control flies (Figure 3C). This was corroborated by an increase in colony forming units (CFUs) in cultured gut extracts (Figure 3D). In addition, 16S ribosomal RNA-sequencing demonstrated clear differences in the intestinal microbes of *Gba1b*^*-/-*^ flies compared to control flies (Figure S4A). In particular, there was a significant increase in the *Acetobacter* and Lactobacillus genera in the gut of *Gba1b*^*-/-*^ flies.

Since *Gba1b*^*-/-*^ flies exhibit an increased bacterial load of altered composition we wished to assess if bacterial challenge was a contributor to early mortality in *Gba1b*^*-/-*^. We orally challenged *Gba1b*^*-/-*^ *flies* with the commensal *Lactobacillus plantarum* bacteria cultured from colonies isolated from control gut microbiota. Infected *Gba1b*^*-/-*^ flies displayed significantly greater mortality than control flies on day 3 post-infection (Figure 3E). Quantitative CFUs assessment showed that there were no significant differences in the *Lactobacillus plantarum* gut load between *Gba1b*^*-/-*^ flies and control after oral infection (Figure S4B). To further understand if the early mortality was due to bacterial proliferation or an overactivation of the innate immune response in the mutant flies, the expression levels of *Drs* were quantified following an intrathoracic injection with heat-killed *Staphylococcus aureus* (*S. aureus*). Aged *Gba1b*^*-/-*^ flies exhibited a ∼3-fold change in *Drs* production compared to controls at 96 hours post injection (Figure S4C). Furthermore, intrathoracic injection with heat-killed *S. aureus* resulted in a reduction in the lifespan of *Gba1b*^*-/-*^ flies compared to flies injected with PBS vehicle alone (Figure S4D). The lifespan of control flies injected with heat-killed bacteria was not significantly different to that of PBS injected flies (Figure S4D). This suggests that it is the inflammatory stimulus and its uncontrolled immune response, rather than bacterial proliferation, that results in compromised survival in *Gba1b*^*-/-*^ flies. Together our data suggest that *Gba1b*^*-/-*^ flies are unable to effectively regulate and respond to commensal gut microbiota, contributing to adverse health outcomes.

### Amelioration of the gut microbiome under germ-free conditions partially rescues survival and locomotor phenotypes in *Gba1b*^*-/-*^ flies

To test whether modulating the microbiome would have beneficial effects, *Gba1b*^*-/-*^ and control flies were reared under germ-free (GF) conditions and behavioural phenotypes were assessed. Adult control and mutant flies were raised and aged on standard food containing a cocktail of antibiotics to maintain GF conditions (Figure 4A). Raising *Gba1b*^*-/-*^ flies under GF conditions resulted in significant reduction of *Dipt* expression and lifespan extension (Figure 4B and 4C). These survival benefits appeared to be specific to *Gba1b*^*-/-*^ flies, as GF control flies did not exhibit reduced AMP expression or improved lifespan compared with non-GF controls. GF conditions also improved locomotor ability in aged *Gba1b*^*-/-*^ flies compared to their non-GF counterparts (Figure 4D). Thus, modulation of the intestinal microbiome partially ameliorates lifespan and locomotor phenotypes in GCase-deficient flies.

**Figure 4.**
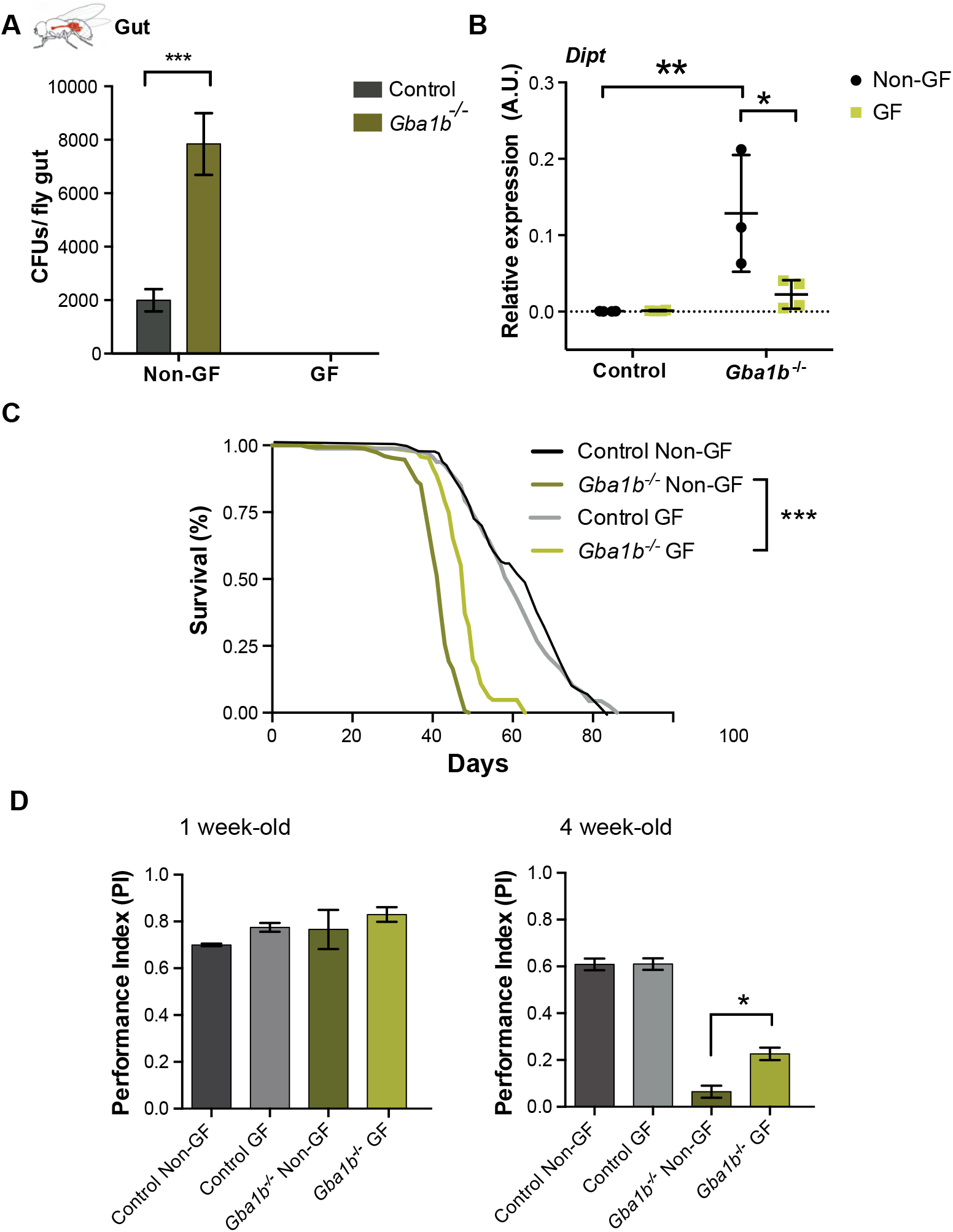
Raising *Gba1* mutant flies under germ-free (GF) conditions partially ameliorates a number of disease phenotypes. **(A)** Colony forming units (CFUs) in *Gba1b*^-/-^ and control fly guts raised under non-GF and GF conditions at 3 weeks. *Gba1b*^-/-^ flies raised under non-GF conditions display higher microbial load in comparison with control flies. No bacterial load is observed for both control and mutant flies raised in GF conditions. Two-way ANOVA test followed by Tukey’s multiple comparison test (***p<0.0001). Data presented as mean ± SD, n=5 per condition. **(B)** Quantitative RT-PCR analysis of *Dipt* mRNA levels in the gut of non-GF and GF *Gba1b*^-/-^ and control flies. *Dipt* levels are significantly reduced in *Gba1b*^-/-^ flies raised in GF conditions. Two-way ANOVA test followed by Sidak’s multiple comparison test (**p=0.0027; *p=0.0107). Data presented as mean ± SD, n=3-4 per condition. **(C)** GF *Gba1b*^-/-^ treated flies have an increased lifespan compared to those reared under standard non-GF conditions. Log-rank tests were used for all comparisons: GF vs non-GF *Gba1b*^-/-^, p=0.0003 (n=150); GF control vs non-GF control p=0.9813 (n=150). **(D)** GF *Gba1b*^-/-^ have improved climbing ability at 4 weeks of age compared to their non-GF counterparts (ns > 0.05; *p=0.034 GF vs non-GF *Gba1b*^-/-^). One-way ANOVA test with Tukey’s post hoc analysis (n=75 flies per condition).

### The gut microbiota regulates the systemic innate immune response in *Gba1b*^*-/-*^ flies

The effect of eliminating the intestinal microbiome on immune activation in the fat body and head was also examined. *Gba1b*^*-/-*^ flies exhibited a striking reduction in fat body Dipt-LacZ staining towards control levels when raised in GF conditions (Figure 5A). Additionally, raising *Gba1b*^*-/-*^ flies under GF conditions resulted in decreased glial activation, as evidenced by a reduction in Draper protein levels in the head (Figure 5B). Remarkably, GF *Gba1b*^*-/-*^ flies displayed a significant reduction of *Dipt* gene expression in the heads to control levels, indicative of reduced IMD activation (Figure 5C). Together, our data suggest that the gut microbiota is responsible for immune activation in the gut and other tissues of GCase-deficient flies and that reversal of this microbiome-dependent immune activation is associated with partial amelioration of lifespan and neuromuscular phenotypes.

**Figure 5.**
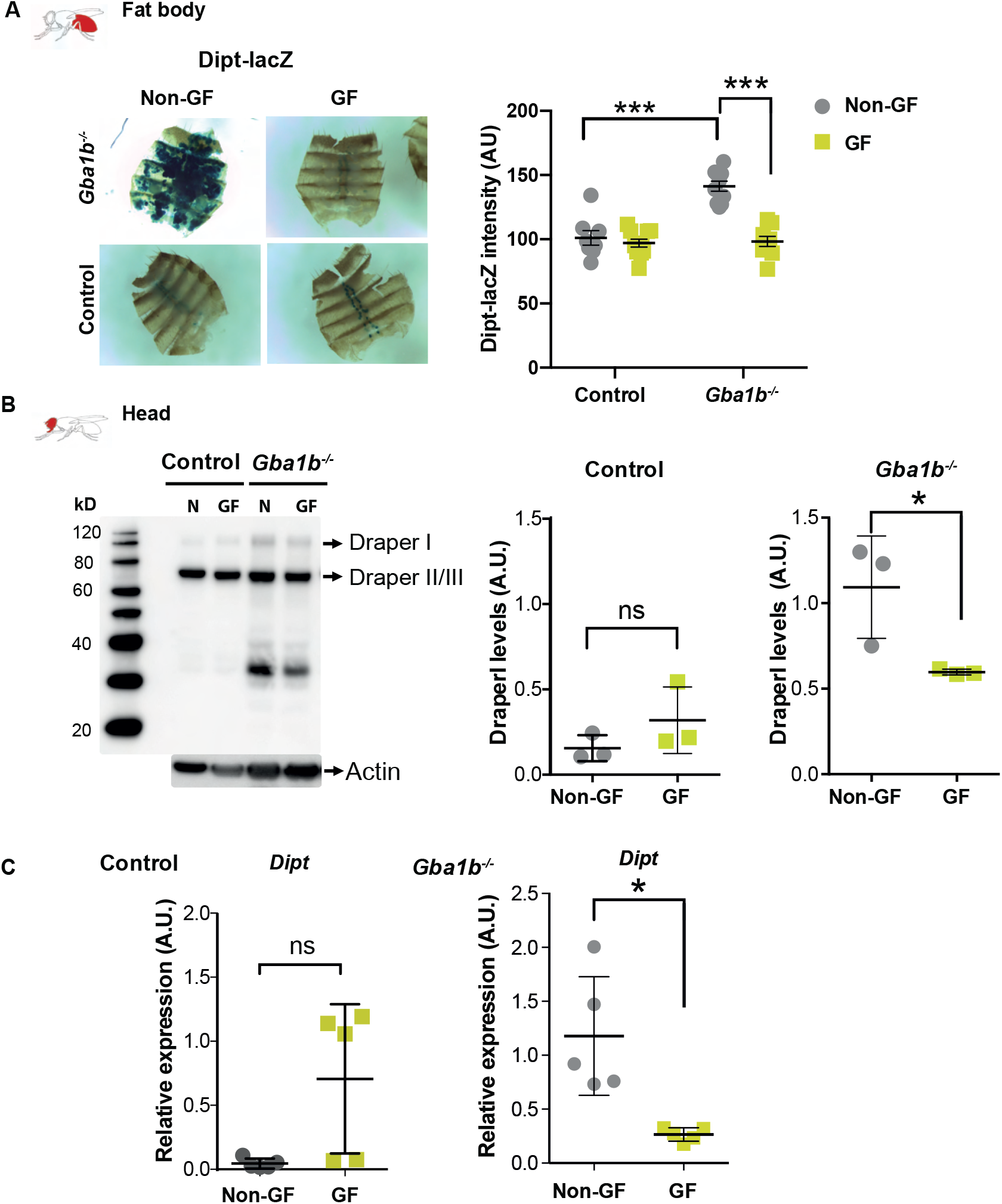
Raising *Gba1b*^*-/-*^ flies under GF conditions reverses immune activation in fat body and brain tissue. **(A)** Dipt-LacZ staining in the fat body is reduced to control levels in GF *Gba1b*^*-/-*^ flies compared to their non-GF counterparts (***p<0.0001). All data was analysed by One-way ANOVA and Tukey’s post hoc analysis and presented as mean ± 95% confidence intervals. **(B)** Western blot analysis reveals a decrease in Draper I levels in the heads of GF *Gba1b*^*-/-*^ flies compared to non-GF flies (n=3; GF *Gba1b*^*-/-*^ vs non-GF *p=0.041; GF controls vs non-GF p=0.991). **(C)** *Dipt* transcript levels are reduced in quantitative RT-PCR analysis from heads of 3-week-old GF *Gba1b*^*-/-*^ flies compared to non-GF flies (n=5 per condition; *p=0.05; unpaired t-test). Data are presented as mean ± SD.

### *Gba1b*^-/-^ flies exhibit blocked autophagy

Autophagy has been shown to have a role in dampening innate immune responses and is achieved by removing key signal transduction components from these pathways (Nakahira et al., 2011, Prabakaran et al., 2018, Tsapras et al., 2022, Tusco et al., 2017). Loss of *Gba1b* has previously been shown to disrupt autophagic flux in the brain in an age-dependent manner (Kinghorn et al., 2016). As the nexus point of microbe-host interactions, autophagy was assessed in the gut of *Gba1b*^*-/-*^ flies. For this purpose the accumulation of Ref(2)P, the fly ortholog of the selective autophagosomal receptor P62, and Atg8a, the fly ortholog of LC3, were used as markers of selective autophagy and macroautophagy respectively (Mauvezin et al., 2014). Western blot analysis on dissected fly guts revealed that both Atg8a-II and Ref(2)P protein levels were significantly increased in *Gba1b*^*-/-*^ flies when compared with control flies (Figure 6A). A significant increase in number and size of Ref(2)P and ubiquitin (Ub) puncta was seen by immunostaining in *Gba1b*^*-/-*^ compared to control guts (Figure 6B). We observed several distinct Lysotracker-positive puncta within the midgut region of 3-week-old *Gba1b*^*-/-*^ flies, but not in guts of control flies (Figure 6C), which is characteristic of impaired autophagic turnover (Kirkin et al., 2009, Nezis et al., 2008). Overall, our data mirrors the autophagy defects seen in the brains of *Gba1b*^*-/-*^ flies (Kinghorn et al 2016).

**Figure 6.**
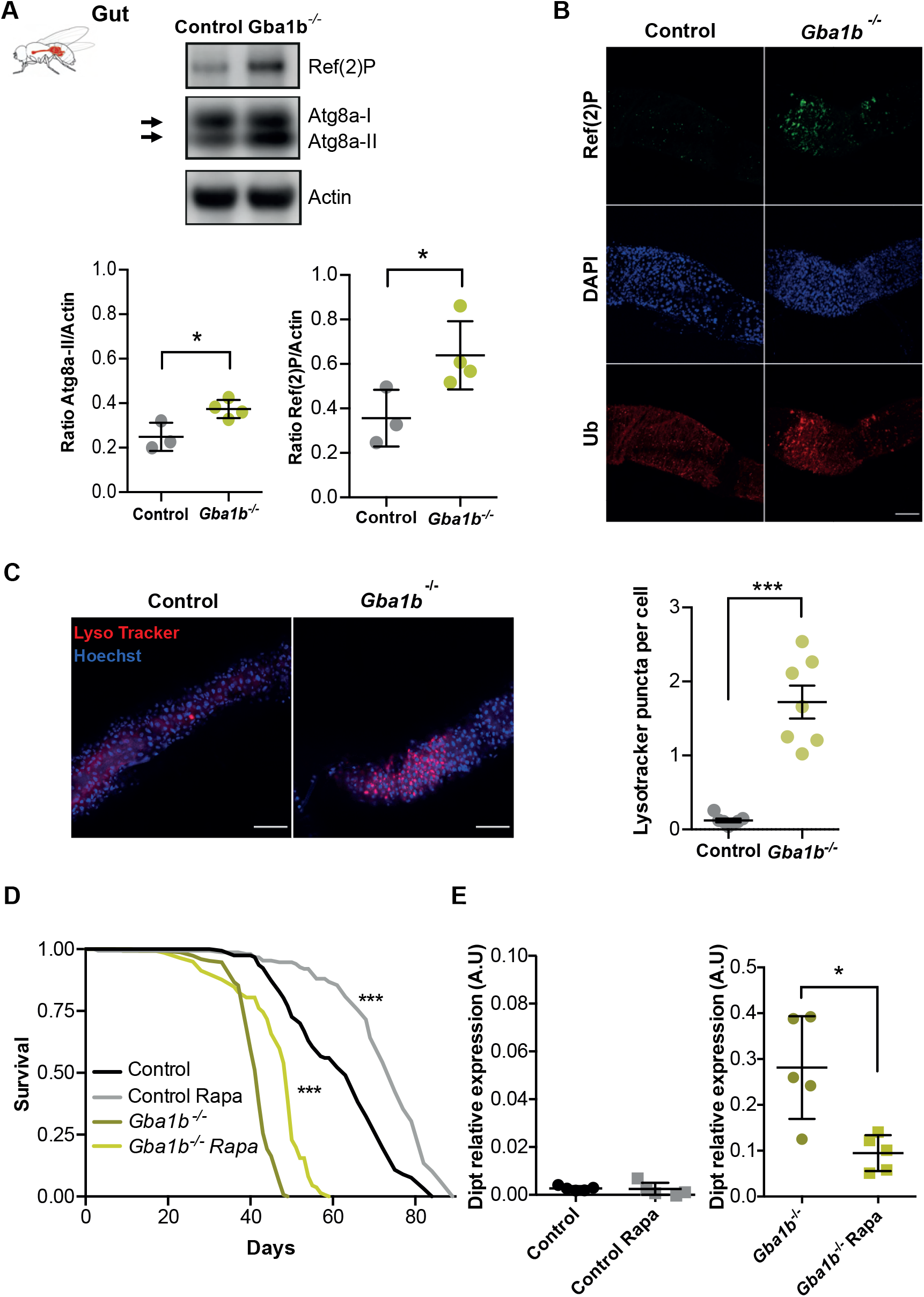
Loss of *Gba1b* results in gut autophagy impairment and stimulation of autophagy through rapamycin administration rescues *Gba1b*^*-/-*^ immune phenotypes and survival. **(A)** Western blot analysis of autophagy-related proteins, Atg8a-I, Atg8a-II and Ref(2)P in fly guts. *Gba1b*^*-/-*^ guts show significant accumulation of Atg8a-II and Ref(2)P proteins relative to control guts (*p=0.024 and *p=0.049, respectively; unpaired t test; n=3-4). Data are presented as mean ± SD. **(B)** Immunostainings of guts labelled with Ref(2)p (green), DAPI (blue) and Ubiquitin (Ub, red). *Gba1b*^*-/-*^ flies display higher number of aggregates of Ref(2)P and ubiquitinated proteins in the middle gut. Scale bar 100 µm. **(C)** Gut immunostaining with Lysotracker (red) and Hoechst (blue) of 3-week-old flies. *Gba1b*^*-/-*^ display an increased number of Lysotracker puncta (***p<0.0001; unpaired t-test). Scale bar 50 µm. **(D)** Lifespan of flies treated chronically with rapamycin (Rapa)(n=150). Survival of treated control and *Gba1b*^*-/-*^ flies is significantly increased (Log-rank tests were used for all comparisons: *Gba1b*^-/-^ vs *Gba1b*^-/-^ Rapa, *** p<0.0001; Control vs Control Rapa *** p<0.0001). **(E)** *Dipt* transcript level is reduced on quantitative q-RT-PCR analysis from guts of 3-week-old *Gba1b*^*-/-*^ flies treated with Rapa compared to non-treated flies (n=5 per condition; *p=0.0159; Mann Whitney test). No significant differences are found in control flies (ns=0.5317). Data are presented as mean ± SD.

### Rapamycin treatment leads to reduced IMD innate immune response and extended lifespan

We hypothesised that the abnormal immune activation that we observe may result from a loss of autophagy-mediated regulation of immune pathways. To investigate this further, we induced autophagy in *Gba1b*^*-/-*^ flies by raising them on rapamycin-supplemented food, a well-known inhibitor of Torc1 and autophagy inducer. Chronic rapamycin treatment resulted in increased levels of Lysotracker-stained puncta in both control and mutant guts and higher ratios of Atg8a II/I indicative of increased autophagic flux (Figure S5). Importantly, rapamycin treatment significantly reduced the levels of the IMD pathway reporter *Dipt* in the gut and significantly extended the lifespan of *Gba1b*^*-/-*^ flies (Figure 6D and 6E).

### Rapamycin treatment efficacy is not enhanced in germ-free *Gba1b*^*-/-*^ flies

Rapamycin treatment has been shown to reduce bacterial load in aged flies (Fan et al., 2015, Schinaman et al., 2019), raising the possibility that the rapamycin-mediated lifespan improvement in *Gba1b*^*-/-*^ was dependent upon the presence of the intestinal microbiota. To study if this was the case, we measured the bacterial load in the guts of treated and non-treated flies with rapamycin by performing quantitative RT-PCR of the 16S rRNA gene and measuring bacteria CFUs (colony formation units) number. Rapamycin treated *Gba1b*^*-/-*^ and control flies exhibited similar expression of 16S rRNA and number of CFUs to non-treated flies. This demonstrates that rapamycin is neither directly bactericidal, nor is its effect as an IMD pathway suppressor contingent on a decrease in bacteria (Figure 7A and B).

**Figure 7.**
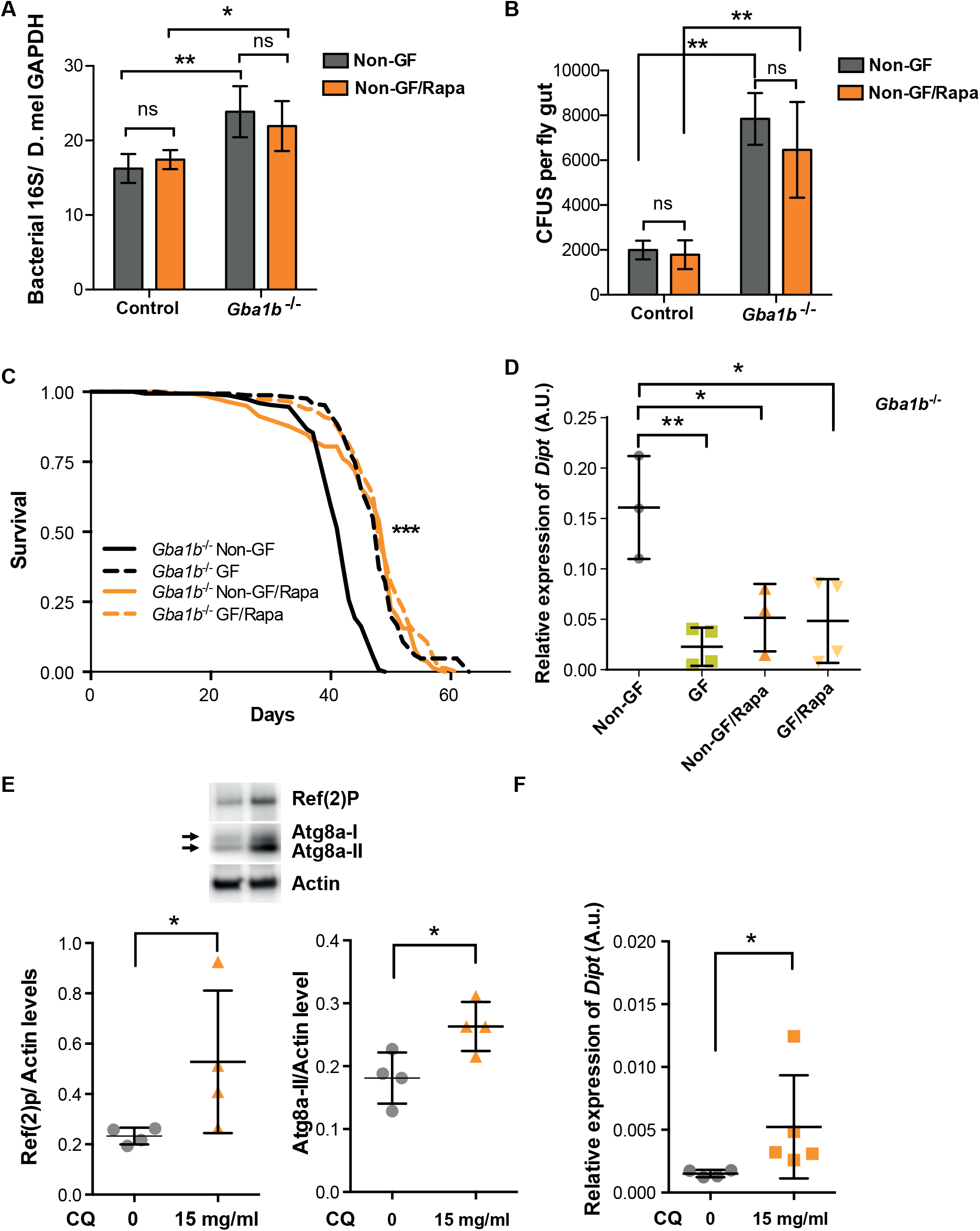
Long-term rapamycin treatment improves *Gba1b*^*-/-*^ survival by reducing inflammation without altering microbial load. **(A)** Quantitative RT-PCR-based 16S rRNA gene from guts of 3-week-old flies does not reveal significant differences in bacterial load of rapamycin (Rapa) treated flies for the two genotypes. Significant differences in bacterial load are observed in Control vs *Gba1b*^*-/-*^ non-GF flies (**p=0.0077) and Control vs *Gba1b*^*-/-*^ non-GF flies treated with Rapa (*p=0.045). Significance was assessed by two-way ANOVA followed by Tukey’s multiple comparison test (n=4 per condition). Colony-forming units (CFUs) assay performed to determine the number of live bacteria from 3-week-old guts of non-GF flies treated or not treated with Rapa. Two-way ANOVA followed by Tukey’s multiple comparison test revealed no significant differences in bacterial load within fly genotype after treatment with Rapa (Control ns=0.991; *Gba1b*^*-/-*^ ns=0.57). Significant differences were found in the comparisons Control vs *Gba1b*^*-/-*^ non-GF flies (**p=0.0022) and Control vs *Gba1b*^*-/-*^ non-GF flies treated with Rapa (**p=0.0087). Data presented as mean ± SD. **(C)** GF and Rapa individual treatments equally extended lifespan of *Gba1b*^*-/-*^ flies (***p<0.0001; log rank test; n=150). No significant additive lifespan extension was observed in *Gba1b*^*-/-*^ flies raised under GF conditions and treated with Rapa. **(D)** Quantitative RT-PCR analysis of *Dipt* transcript levels in guts from *Gba1b*^*-/-*^ flies raised in GF, Rapa and GF/Rapa conditions. One-way ANOVA followed by Tukey’s multiple comparison test (non-GF vs GF, **p=0.0030; non-GF vs non-GF/Rapa, *p=0.0207; non-GF vs GF/Rapa, *p=0.0117; GF vs non-GF/Rapa, p=0.7434; GF vs GF/Rapa, p=0.766; non-GF/Rapa vs GF/Rapa, p=0.999). Data presented as mean ± SD (n=3-4). **(E)** Western blot analysis of Ref(2)P and Atg8a on guts from 3-week-old control flies treated with chloroquine (CQ) for 48 hours. CQ treatment significantly induces accumulation of Ref(2) (*p=0.028; Mann Whitney test; n=4) and Atg8a-II (*p=0.0269; n=4). **(F)** *Dipt* transcript level is increased on quantitative RT-PCR analysis from guts of 3-week-old control flies treated with CQ for 48 hours compared to non-treated flies (*p= 0.0159; Mann Whitney test; n=4 per condition). Data are presented as mean ± SD.

To directly determine whether rapamycin and GF treatment exert non-overlapping independent effects, GF and non-GF *Gba1b*^-/-^ flies were treated with rapamycin or vehicle control (EtOH). Rapamycin similarly extended the lifespan of mutants under GF and non-GF conditions (Figure 7C). No additive effect was observed when flies were treated with rapamycin in the absence of the microbiota, suggesting both treatments are partially rescuing the fly survival through a common downstream mechanism. As rapamycin treatment reduces *Dipt* levels, we tested whether *Gba1b*^-/-^ GF flies treated with rapamycin express further lower levels of this AMP. In agreement with the effect on lifespan, we found that GF mutant flies treated with rapamycin had similar levels of *Dipt* when compared with non-GF *Gba1b*^-/-^ flies treated with rapamycin (Figure 7D). Together, our results indicate that reduction of the innate immune response either by improving intestinal autophagic degradation of putative immune cascade components or elimination of the microbiota is sufficient to improve lifespan of the *Gba1b*^*-/-*^ flies.

To further substantiate the link between defective intestinal autophagy and persistent activation of the innate immune response, we inhibited late-stage autophagy in control flies. For that purpose, we fed flies with chloroquine (CQ), a well-characterized autophagy inhibitor that blocks autolysosome acidification and fusion, then measured the effect on the autophagic and immune pathways. Feeding flies with CQ resulted in accumulation of Atg8a-II and Ref(2)P (Figure 7E) and significantly increased *Dipt* expression in the gut (Figure 7F).

Overall, these data demonstrate that basal autophagy is intricately involved in control of the innate immune response and that blockage of autophagy in the gut is sufficient to recapitulate immune phenotypes seen in *Gba1b*^-/-^ fly guts.

## Discussion

The immune system and gut-brain axis communication are increasingly being implicated in neurodegenerative disease. Using a GCase loss-of-function fly model of GD, we demonstrated that an altered gut microbiota, coupled with intestinal autophagic dysfunction, directly stimulates innate immune activation, which is detrimental to organismal health.

Our genome-wide genetic analysis revealed that loss of GCase activity leads to an age-dependent increase in the expression of Toll, IMD and JAK-STAT pathway components. The *GBA1* gene is widely expressed in the human brain (proteinatlas.org) (Uhlen et al., 2015) and shows significantly higher expression in astrocytes and microglial/macrophage cells compared to neurons (brainrnaseq.org). Moreover, here we showed increased Draper-dependent glial activation in the *Gba1b*^*-/-*^ fly brain. Microglia, the dominant innate immune cells of the mammalian brain, are increasingly being implicated in neurodegenerative disease (Hickman et al., 2018). Reactive microglia are present in post-mortem PD brains (McGeer et al., 1988) and positron emission studies have similarly reported increased microglial responses in cortical and subcortical areas in vivo in early PD, as well as in the brains of *GBA1* mutation carriers without PD (Gerhard et al., 2006, Mullin et al., 2021, Terada et al., 2016). Mouse models of neuronopathic GD have also revealed widespread astrogliosis and microgliosis, in association with fatal neurodegeneration within the first weeks of life (Farfel-Becker et al., 2011, Massaro et al., 2018). In addition to innate immune activation in the fly head, we confirmed that *Gba1b*^*-/-*^ flies display elevated peripheral innate immune responses, including in the gut and fat body. This is in keeping with a previous study in flies harbouring insertions of minos elements in the two *Gba1* genes, resulting in the production of truncated mutant GCase proteins. These flies displayed up-regulation of IMD and Toll pathway genes in the head and body (Cabasso et al., 2019, Cabasso et al., 2021). Our results thus confirm that complete loss of *Gba1b* gene activity is associated with strong innate immune activation, including the JAK-STAT stress response.

Our finding that innate immune activation occurs peripherally is consistent with studies showing evidence of significantly higher levels of serum proinflammatory cytokines and altered immune cell profiles in the blood of PD and GD patients compared to heathy individuals (Barak et al., 1999, Williams-Gray et al., 2016). There is increasing evidence for intestinal dysfunction in PD (Devos et al., 2013, Schwiertz et al., 2018), but it is not known whether gut pathology is a feature of *GBA1*-PD or GD. We demonstrated gut dysfunction with increased intestinal transit time and elevated gut wall permeability in *Gba1b*^*-/-*^ flies. Further probing revealed increased innate immune activation, predominantly of the IMD and Toll pathways, in the intestinal tissue of *Gba1b*^*-/-*^ flies. It is likely that these gut pathologies are a consequence of GCase loss-of-function in the intestinal tissue, as we demonstrated the *Gba1b* gene is expressed in the *Drosophila* gut. Importantly, our findings reflect those seen in PD patients where enteric inflammation on colonic biopsies has been reported, including increased intestinal expression of pro-inflammatory cytokines such as TNF-a, IL6 and IL-1b, and the bacterial endotoxin ligand TLR4 (Devos et al., 2013, Perez-Pardo et al., 2019). Furthermore, intestinal permeability is increased in PD patients in association with decreased serum LPS binding protein (LBP), indicating greater endotoxin exposure (Forsyth et al., 2011, Pal et al., 2015). The intestinal microbiota is essential for the management of gut epithelial integrity, through maintenance of tight junction proteins and mucin production, thereby inhibiting the infiltration of bacteria and immunogenic products (Michie and Tucker, 2019).

The *Drosophila* midgut is an ideal model system for studying host-microbiome interactions due to striking conservation of intestinal structure and gut immune signalling with humans (Liu et al., 2017). Consistent with the intestinal microbiome playing a role in *GBA1*-associated neurodegeneration, we observed increased intestinal bacterial load and alterations in the composition of the gut microbiome in *Gba1b*^*-/-*^ flies. In particular, there was a significant increase in the abundance of *Acetobacter* and *Lactobacillus*, the commonest genera in the fly gut (Heine et al., 2001). The finding that bacterial load was increased in *Gba1b*^*-/-*^ flies compared to control flies is commensurate with the observation that bacterial load is a significant determinant of lifespan in *Drosophila* (Lee et al., 2019). Our results also mirror several studies reporting changes in a number of gut bacterial species between PD patients and control groups (Caputi and Giron, 2018, Lin et al., 2019, Vidal-Martinez et al., 2020).

The chronic gut immune activation observed in *Gba1b*^*-/-*^ flies likely promotes further microbiome dysbiosis, which in turn can lead to impairment of gut function and ultimately reduced survival (Clark et al., 2015). Support for intestinal microbiome-mediated effects on the CNS in *Gba1*^-/-^flies, i.e. via gut-brain axis communication, was achieved by raising mutant and control flies under germ-free (GF) conditions. Modulation of the intestinal microbiome in GF flies resulted in partial amelioration of a number of phenotypes, including glial activation and locomotor defects. Similar beneficial effects were seen on modulation of the intestinal microbiota in Alzheimer’s disease mouse and fly models (Harach et al., 2017, Kim et al., 2020, Westfall et al., 2019) and α-Syn overexpressing (ASO) mice (Sampson et al., 2016). In the case of ASO mice, amelioration of the intestinal microbiome led to an improvement in neuropathology, whereas oral administration of specific microbial metabolites to GF mice promoted neuroinflammation and motor deficits. Interestingly, we show that the re-introduction of a single commensal bacteria from the gut of healthy control flies is sufficient to promote immune activation in *Gba1b*^*-/-*^ flies in association with significant mortality.

The finding that the intestinal microbiome in *Gba1b*^*-/-*^ flies is necessary to promote innate immune activation can be interpreted in the context of the recently described endotoxin hypothesis of neurodegeneration (Brown, 2019). Endotoxins predominantly refer to the lipopolysaccharide found in the outer cell wall of gram-negative bacteria (Heine et al., 2001). We hypothesise that the intestinal inflammation and altered gut microbiome observed in our *Gba1b*^*-/-*^ fly model promotes intestinal barrier permeability (‘gut leakiness’). This leads to translocation of microbiota-shed pathogen associated molecular patterns (PAMPs) -among them endotoxins and peptidoglycan fragments-from the gut lumen into the systemic circulation (Brown, 2019, Pal et al., 2015). The increased intestinal transit time and increased bacterial load observed in *Gba1b*^*-/-*^ flies may further promote this translocation due to the increased exposure of the gut wall to bacteria. The subsequent circulating microbial products can stimulate a chronic systemic innate immune activation produced by fat body (Duerkop et al., 2009), resulting in breakdown of the blood brain barrier and neuroinflammation (Fukui, 2016, Logsdon et al., 2018, Perez-Pardo et al., 2019).

Fedele and colleagues recently showed that in *pink-1* mutant flies display increased Relish NF-kB signalling with concomitant intestinal disfunction and cell death in the midgut. Intriguingly, the neuronal defects observed in these flies can be rescued by genetic inhibition of Relish transcription factor (Fedele et al., 2022).

Furthermore, we showed that blockage of autophagy in healthy flies is sufficient to induce an immune response in the gut. This is in agreement with recent studies that show degradation of IMD pathway mediators, such as Tak1/Tab2 and IKK complex is mediated by selective autophagy. Loss of normal autophagic flux results in constitutive activation of the IMD pathway (Tsapras et al., 2022, Tusco et al., 2017). Autophagy is significantly impaired in the brain, gut and fat body tissue of *Gba1b*^*-/-*^ flies. This in turn may lead to an inability to terminate an activated innate immune response, through the degradation of key components of the NF-*K*B signalling cascades, resulting in increased production of AMPs. We therefore propose that the deleterious chronic inflammation observed in *Gba1b*^*-/-*^ flies results from the loss of autophagic regulation of immune signal transduction pathways.

In conclusion, we demonstrate innate immune activation, GI dysfunction, and microbiome dysbiosis in a fly model of GCase loss-of-function. We show that the intestinal microbiota stimulates local gut and systemic innate immune responses in *Gba1b*^*-/-*^ flies. We also demonstrate that reduction of the innate immune response either by removal of the microbiome or improvement of autophagy improves survival as well as gut and locomotor phenotypes. This insight has the potential to lead to the development of novel long-awaited therapeutic strategies in the treatment of PD and neuronopathic GD.

## Material and Methods

### Drosophila stocks and handling

All flies were backcrossed at least 6 generations into the *w*^1118^ background to create isogenic background lines. Flies lacking *Gba1b* (*Gba1b*^*-/-*^) were previously described (Kinghorn et al., 2016). All fly stocks were maintained at 25°C on a 12:12 hour light: dark cycle at constant humidity on a standard sugar-yeast medium (15 g/L agar, 50 g/L sugar, 100 g/L autolysed yeast 3 g/L nipagin and 3 ml/L propanoic acid). *Gba1b* knockout flies were kept in mixed populations with control flies for several generations prior to setting up all experiments, except for those involving the reintroduction of *Lactobacillus plantarum*. This was performed to reduce the confounding effects of environmental conditions.

The *Gba1b* CRIMIC Trojan-Gal4 line (TI(CRIMIC.TG4.0}Gba1bCR00541-TG4.0; stock number 78943) was obtained from the Bloomington Stock Centre (Indiana, USA). All experiments were set up using ‘egg squirt protocols’ to ensure that all experimental flies were raised at similar larval densities (∼300 eggs per bottle containing 70 mL of food). Briefly, flies were allowed to lay eggs for less than 24 hours on grape medium plates, with live yeast paste to encourage egg-laying. The eggs were collected from the plate by washing with phosphate buffered saline (PBS) solution and collected into falcon tubes. The eggs were then allowed to settle to the bottom of the tube. Using a 100 μL Gilson pipette, ∼18-21 µL of egg suspension was dispensed into 200 mL glass 16 bottles containing 70 mL SY medium. Following eclosion, flies were transferred to fresh food for a 48-hour mating period. Under CO2 anaesthesia, flies were then divided into 15 female flies per vial.

### RNA sequencing analysis

Next generation sequencing was performed by the Glasgow Polyomics (UK) facility. Using our in-house protocol RNA was isolated from the *Gba1b*^*-/-*^ heads and the heads of aged-matched *w*^1118^ controls at 1 week and 3 weeks-old of age. For each sample time point, five replicates of 30 flies were used. The twenty samples were PolyA-enriched and the libraries were sequenced on a MiSeq Illumina instrument as 75 bp paired-end reads generating a total of 25 million reads. The mean output per sample was 10 million reads. Reads were mapped to Drosophila melanogaster genome (downloaded from FlyBase.org, version dmel_r6.31_FB2019_06) using STAR v2.7.3a (Dobin et al., 2013) (https://doi.org/10.1093/bioinformatics/bts635) [default parameters], plus the ENCODE options for ‘--alignIntronMin’, ‘--alignIntronMax’ and ‘--outFilterMultimapNmax’. Following reads sorting by gene name using STAR, featureCounts v.2.0.0 (Liao et al., 2014) (https://doi.org/10.1093/bioinformatics/btt656) [default parameters] was used to extract gene counts as per the transcriptome annotations (downloaded from FlyBase.org, version dmel_r6.31_FB2019_06/gtf). Differential expression analysis was performed for each time point independently vs the aged-matched controls using DESeq2 (Love et al., 2014) (https://doi.org/10.1186/s13059-014-0550-8) [default parameters]. The generated p values were adjusted for multiple testing using the procedure of Benjamini and Hochberg (Benjamini., 1995). Gene lists were created by filtering for lowly expressed genes (5 reads across all five replicates) and using absolute cut-offs for *p-value* <0.05 and log2 fold change ≥2 or log2 fold change ≤-2 for up-regulated and down-regulated genes, respectively. Common genes at both time points were obtained by merging together the significant gene lists. These were further filtered by log2 fold change ≥2 at the latter time point generating common genes. Genes were visualised as a heat map created using the ‘heatmply’ R package v.1.1.0. Gene ontology (GO) enrichment analysis was performed using g:Profiler (Raudvere et al., 2019) (https://doi.org/10.1093/nar/gkz369).

### Quantitative RT-PCR

Total RNA was extracted from 6 whole flies, 25 heads, 10 dissected midguts or 10 headless bodies of adult flies at each time point using Trizol (Invitrogen) according to the manufacturer’s instructions. 4 µg of total RNA for each sample was subjected to DNA digestion using DNAse I (Invitrogen), immediately followed by reverse transcription using the Superscript II system (Invitrogen) with oligo(dT) primers. Quantitative RT-PCR was performed using the QuantStudio 6 Flex detection system (Applied Biosystems) and SYBR Green (Invitrogen) as per the manufacturer’s instructions. Each sample was analysed in duplicate, and the values are the mean of at least 4 independent biological repeats. The primers used were as follows:

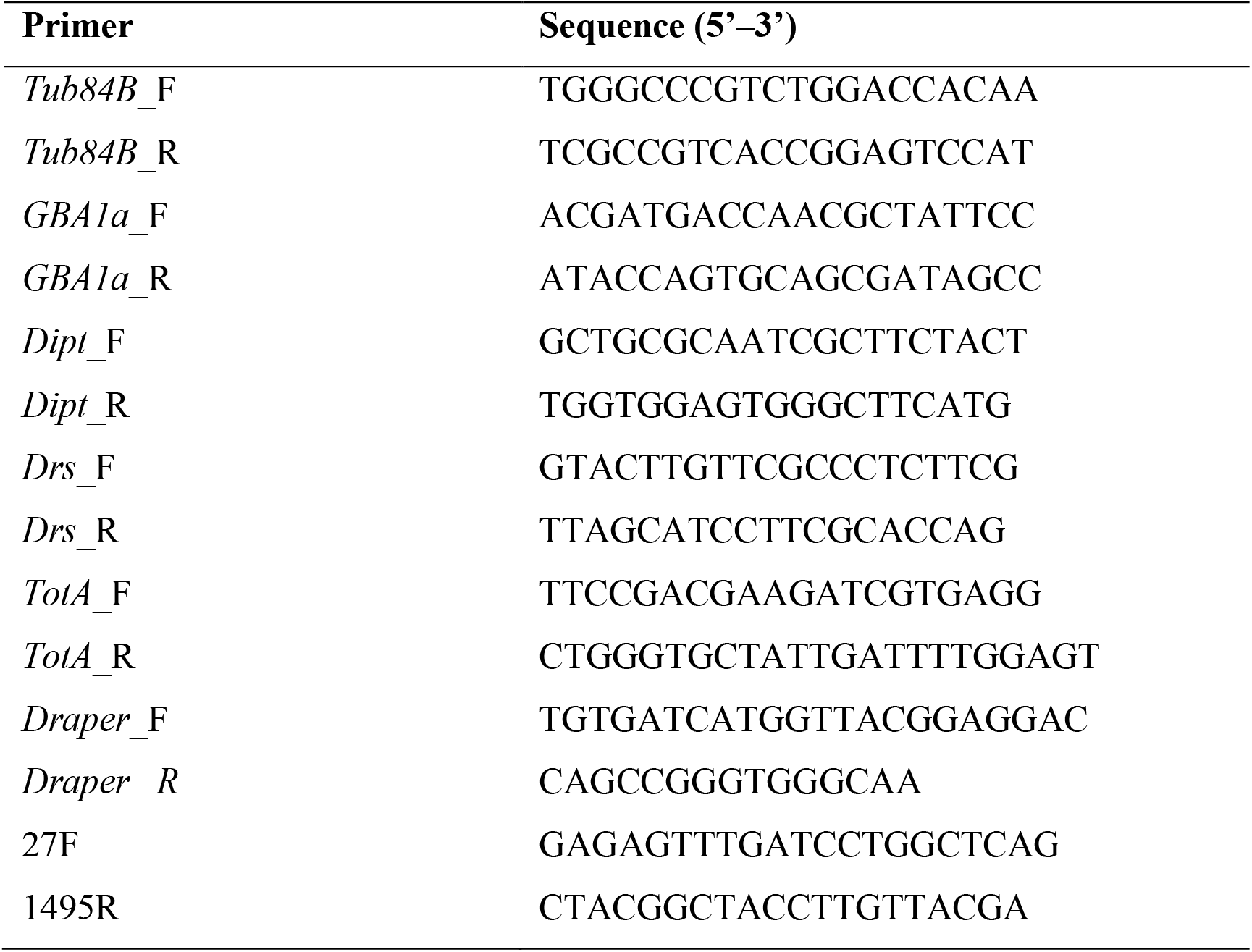

### Longevity and climbing assays

Flies were transferred to vials containing fresh food three times a week throughout life. The number of dead flies found during each transfer was recorded. Lifespan curves were analysed using a log-rank test. For climbing assays, 75 flies of each genotype were housed in groups of 15 in plastic vials held in a Drosoflipper (http://drosoflipper.com/). While video recording, the flies were tapped to the bottom and allowed to climb for 30 seconds. The numbers of flies in the top, middle and bottom thirds of each vial at 30 seconds were scored. A performance index was calculated as previously described (Yu et al., 2020).

### Germ-free (GF) conditions

Flies were rendered GF as previously described (Sabat, 2015). Axenic food was prepared with 50 mg/L tetracycline (Sigma-Aldrich, T3258) and 400 mg/L streptomycin (Sigma-Aldrich, S9137). In brief, embryos were collected on grape juice agar plates for 24 hours and transferred to a nylon basket. Embryos were dechorionated in 50% bleach for 2 minutes, washed twice in 70% EtOH for 1 minute and then washed in dH2O for 10 minutes. Dechorionated embryos were then transferred onto antibiotic food. Flies were processed at 3 weeks of age.

### Western blot analysis

For Western blotting, 10 fly heads or 8 guts per sample were homogenized in 2× Laemmli loading buffer (100 mM Tris 6.8, 20% glycerol, 4% SDS) containing 5% β-mercaptoethanol and then boiled for 5 minutes. Approximately 8 ul of protein extract was loaded per lane. Proteins were separated on precast 4%–12 % NuPage Bis-Tris gels (Invitrogen) and transferred to a PVDF membrane. The membranes were then blocked in 5% BSA in TBST (TBS with 0.05% Tween 20) for 1 hour at room temperature, after which they were probed with primary antibodies diluted in 5% BSA in TBST overnight at 4°C. Blots were developed using the ECL detection system. Primary antibodies used: mouse DHSB Draper 5D14 (1:500); rabbit anti-GABARAP (ab109364, 1:2000); rabbit anti-Ref(2)P (ab178440, 1:500). The secondary antibody (Abcam) was diluted 1:10.000 (goat anti-mouse: ab6789 or goat anti-rabbit: ab672) in 5% BSA in TBST for 1 hour at room temperature. Bands were visualized with Luminata Forte (Millipore). All blots were imaged with ImageQuant LAS4000 (GE Healthcare Life Science). Quantification was performed using the ImageJ program (National Institutes of Health).

### Immunofluorescence microscopy

Tissue dissections were modified from Wu et al. (2006) (Wu and Luo, 2006). Brains, guts or fat bodies were dissected in PBS and fixed, nutating in 4% paraformaldehyde (in PBS) for 20 minutes. Dissected tissues were washed twice with PBS-T (PBS, 0.5% TritonX), followed by 3x 20 minute washes in PBS-T. After, tissues were incubated in block solution (PBS-T with 5% H1 horse serum (Gibco) for 1 hour before transferral to Block containing primary antibody overnight at 4 °C. Tissues were then washed twice in PBS-T, followed by 3x 20 minutes in PBS-T and transferred to block solution containing secondary antibody overnight at 4 °C. Tissues were then washed twice in PBS-T, followed by 3x 20 minutes in PBS-T and mounted in Vectashield antifade mounting medium with DAPI (Vector Labs, H1200). For experiments co-staining alongside mCherry, antibody incubations were reduced to 2 hours at room temperature to preserve mCherry signal. Antibodies working concentrations were as follows: mouse anti-Draper 5D14 (1:500), mouse anti-repo 8D12 (1:200), mouse anti-elav 958A9 (1:200), rabbit anti-Ref(2)P (ab178440, 1:200), mouse anti-FK2 (Sigma 04263, 1: 800). The secondary antibodies used: goat anti-mouse Alexa488 (A11001), and rabbit Alexa568 (A11036) were used at 1:250.

### Lysotracker staining

Fly guts were dissected in PBS and stained with lysotracker (LysoTracker Red DND-99; Thermo Fisher Scientific; 1:2000) and Hoeschst 33342 (Sigma, 1mg/ml; 1:1000) for 3 minutes. Immediately after staining, guts were washed 2x for 5 min with PBS, mounted with Vectashield and imaged.

### Dipt-LacZ staining

Dipt-LacZ expressing abdomens were dissected in ice-cold PBS and immediately fixed in 4% PFA for 20 minutes. Abdomens were washed in PBS and transferred into staining solution (10 mM potassium ferricyanide, 10mM potassium ferrocyanide, 1mM MgCl2, 150mM NaCl, 0.1% Triton in PBS), pre-warmed to 37°C containing 1:20 X-gal (1 mg/ml in DMSO) and incubated overnight in the dark. Abdomens were washed 3x in pre-warmed PBS, mounted and immediately imaged on a Leica M165C light microscope. LacZ intensity was quantified by splitting RGB channels, inverting the image and quantifying the entire abdomen, with the background subtracted.

### Gut transit time and faecal deposit characterisation

Flies were placed on SY diet containing 0.5 % Bromophenol Blue (BPB) (Sigma-Aldrich, B5525-10G) for 36 hours, then transferred into empty glass vials every 2 hours. Faecal deposits were then manually counted at each timepoint and assessed for ROD morphology.

### Gut permeability analysis (Smurf assay)

Gut barrier efficiency was analysed by placing flies on blue food prepared using 2.5% (w/v) FD°C blue dye n° 1 (Fastcolors) as previously described (Regan et al., 2016), aged flies were kept on blue food for 6 days and subsequently scored for Smurf phenotype.

### Image analysis

All images were acquired on a Zeiss LSM 700 confocal microscope and all settings were kept the same within an experiment. For Draper staining, a maximum intensity 10 µm z-stack with 1µm intervals encompassing the antennal lobes was taken. Regions of interest, encompassing the soma, were generated using the DAPI channel as a guide. Intensity relative to the antennal lobe neuropil was measured.

### Preparation of template DNA for 16S ribosomal RNA sequencing on fly midgut tissue

Extracted midguts were added to an Eppendorf tube containing 180 μL Lysis buffer (20 mM Tris-HCl pH 8.0, 2 mM EDTA pH 8.0, 1.2 % Triton X-100 (Sigma-Aldrich), 20mg/mL fresh lysozyme from chicken egg (Sigma-Aldrich L7651)). 200 μL QIAGEN buffer AL was added to each Eppendorf tube containing 40 midguts. Following lysis with a Kontes pellet pestle, 20 μL proteinase K (QIAGEN) was added. The samples were then incubated at 56 °C for 3 h. To remove any RNA, 10 μL of 10 μg/mL RNase A (Sigma-Aldrich R4875) was added and the samples were incubated at 37 °C for 30 minutes. 200 μL ethanol was then added and the standard QIAGEN spin column protocol was followed. The DNA concentration was then assessed using a Nanodrop 2000 Spectrophotometer (Thermo Fisher Scientific) and sent to LC Sciences (Houston, USA) on dry ice for 16S/18S/ITS1/ITS2 RNA-sequencing.

### 16S ribosomal RNA sequencing

The V3-V4 region of the prokaryotic (including bacterial and archaeal) small-subunit (16S) rRNA gene was amplified with slightly modified versions of primers 338F (5’-ACTCCTACGGGAGGCAGCAG-3’) and 806R (5’-GGACTACHVGGGTWTCTAAT-3’). The 5’ ends of the primers were tagged with specific barcodes and sequencing universal primers. PCR amplification was performed in 25 μL of reactions containing 25 ng of template DNA, 12.5 μL of PCR premix, 2.5 μL of each primer, and PCR-grade water. The PCR conditions for amplifying the prokaryotic 16S fragments comprise the following steps: initial denaturation at 98 °C for 30 seconds; 35 cycles of denaturation at 98 °C for 10 seconds, annealing at 54 °C/52 °C for 30 seconds, and extension at 72 °C for 45 seconds; and final extension at 72 °C for 10 minutes. PCR products were confirmed with electrophoresis in 2 % agarose gel. Ultrapure water was used as the negative control to exclude false positives. PCR products were purified by AMPure XT beads (Beckman Coulter Genomics, Danvers, MA, USA) and quantified by Qubit (Invitrogen, USA). The size and quantity of the amplicon library were assessed with Agilent 2100 Bioanalyzer (Agilent, USA) and Library Quantification Kit for Illumina (Kapa Biosciences, Woburn, MA, USA), respectively. PhiX control library (v3) (Illumina) was combined with the amplicon library (at a fraction of 30 %). The libraries were sequenced on Illumina MiSeq (300 bp ’ 2, pair-ended) using the standard Illumina sequencing primers.

### 16S ribosomal RNA sequencing data analysis

Paired-end reads were assigned to samples based on their unique barcodes before barcode and primer sequences were trimmed. The trimmed reads were merged using FLASH. Quality filtering on the raw tags were performed to obtain high-quality clean tags with *fqtrim* (v0.94). Chimeric sequences were filtered using *Vsearch* (v2.3.4). Sequences with ≥97% similarity were assigned to the same operational taxonomic units (OTUs) by *Vsearch* (v2.3.4). Representative sequences were chosen for each OUT, followed by taxonomic assignment using the RDP (Ribosomal Database Project) classifier. The differences of the dominant species in different groups and multiple sequence alignment were conducted by *mafft* software (v7.310). OTU abundance information were estimated after rarefaction with the least sequence number obtained for the project. Alpha diversity is applied for analyzing complexity of species diversity with 5 measurements, including Chao1, Observed species, Goods_coverage, Shannon, Simpson, which were calculated by QIIME (v1.8.0). Beta diversity was calculated by PCoA analysis to evaluate differences of samples in species complexity. Cluster analysis was performed by QIIME software (v1.8.0).

### 16S qPCR quantification of microbiota load

DNA was extracted from samples containing 5 female flies. Each fly was first sterilized with 70 % ethanol to remove exterior bacteria and then washed with 1x PBS. After this step, the same protocol as described above was used to extract the DNA for the 16S Ribosomal RNA gene sequencing. qPCR was then performed on total genomic DNA to determine the bacterial load to fly DNA in each sample by normalizing it to the *Drosophila* GAPDH gene.

### CFUs and analysis of culturable bacteria

Culture-dependent methods were used to quantify and identify culturable bacteria present in the fly gut. Guts were dissected from female flies in Tris-HCl 50 mM, pH 7.5 and homogenized with a Kontes pellet pestle in 300μL Luria-Bertani medium (BD™ Difco). Each sample was then serially diluted and 50 μL from each dilution were plated onto different agar media: LB (BD™ Difco), Brain heart infusion (BHI) (Sigma-Aldrich), de Man, Rogosa and Sharpe broth (MRS) (Thermo Scientific ™) and Mannitol (3 g of Bacto Peptone [Becton Dickinson], 5 g of Yeast Extract [Sigma-Aldrich], 25 g of D-Mannitol [Sigma-Aldrich], 1 L of Milli-Q water). Liver Infusion Broth (Becton Dickinson). Plates were incubated at 28°C for three days and isolated colonies present in each plate were counted using a digital colony counter (Fisher Scientific). Five colonies for each morphological type were re-streaked into new plates. In order to identify each isolate a PCR was performed using part of a colony as DNA template and 16S universal primers (27F and 1495R). The amplified amplicon was cleaned up using PCR purification kit (Qiagen) and sent to Source Biosciences for sequencing. Identified isolates were grown in liquid medium containing 25% of glycerol (v/v) and frozen at -80°C.

### Oral infection with *Lactobacillus plantarum*

Dissected guts from 6 *w*^1118^ control female flies (previously washed in EtOH and PBS) were homogenised and plated onto MRS agar medium and incubated at 25 °C for two days. Colonies were then isolated and identified by 16S Ribosome Gene Sequencing (Source Bioscience). A single colony of *Lactobacillus plantarum* was then grown overnight in 400 ml of MRS broth at 25 °C. The cell culture was centrifuged at 5000 rpm (10 min), washed in PBS and resuspended in 5 ml of 2.5 % of sucrose. 200 μl of this suspension or 2.5% sucrose vehicle was pipetted onto 2 filter papers (Whatman filter paper Chroma circles 2.1 cm) in a vial containing 1.5 % agarose. 3-week-old flies were transferred to vials containing either 2.5 % sucrose and *Lactobacillus* or 2.5 % sucrose alone. After 72 hours of feeding, fly death was scored.

### Micro-injection of flies with heat-killed bacteria

Newly eclosed *Gba1b* mutant and control flies were reared at 25°C on SY food until approximately 3 weeks of age. They were then subjected to an injection into the thorax with 32 nl heat-killed *Staphylococcus aureus* NCTC8325-4 (BAC) or PBS using a microinjector (Nanoject II; from Drummond Scientific).

### Statistical Analyses

Statistical analyses were performed with Prism6 (GraphPad Software, USA). Data were tested for normal distribution and equal variance and accordingly analysed using adequate statistical tests – described in the legend of each figure. Statistical differences were considered significant at p<0.05. Log-rank test on lifespan data were performed in Microsoft Excel (template 351 available at http://piperlab.org/resources/) and data was plotted using Prism 6.

## ACKNOWLEDGEMENTS

We thank John Labbadia, Nathaniel Woodling, Daniel Fabian, Matias Fuentealba and all members of the Institute of Healthy Ageing, as well as Hugo Bellen and Liping Wang (Baylor College of Medicine) for their critical input. This work was supported by the Wellcome Trust (K.J.K. (Wellcome Trust Career Development Fellowship) 214589/Z/18/Z), funding from the Rosetrees Trust (K.J.K. and A.H. (M701 and M701A)) and the Academy of Medical Sciences (K.J.K).

---

**Figure S1.**
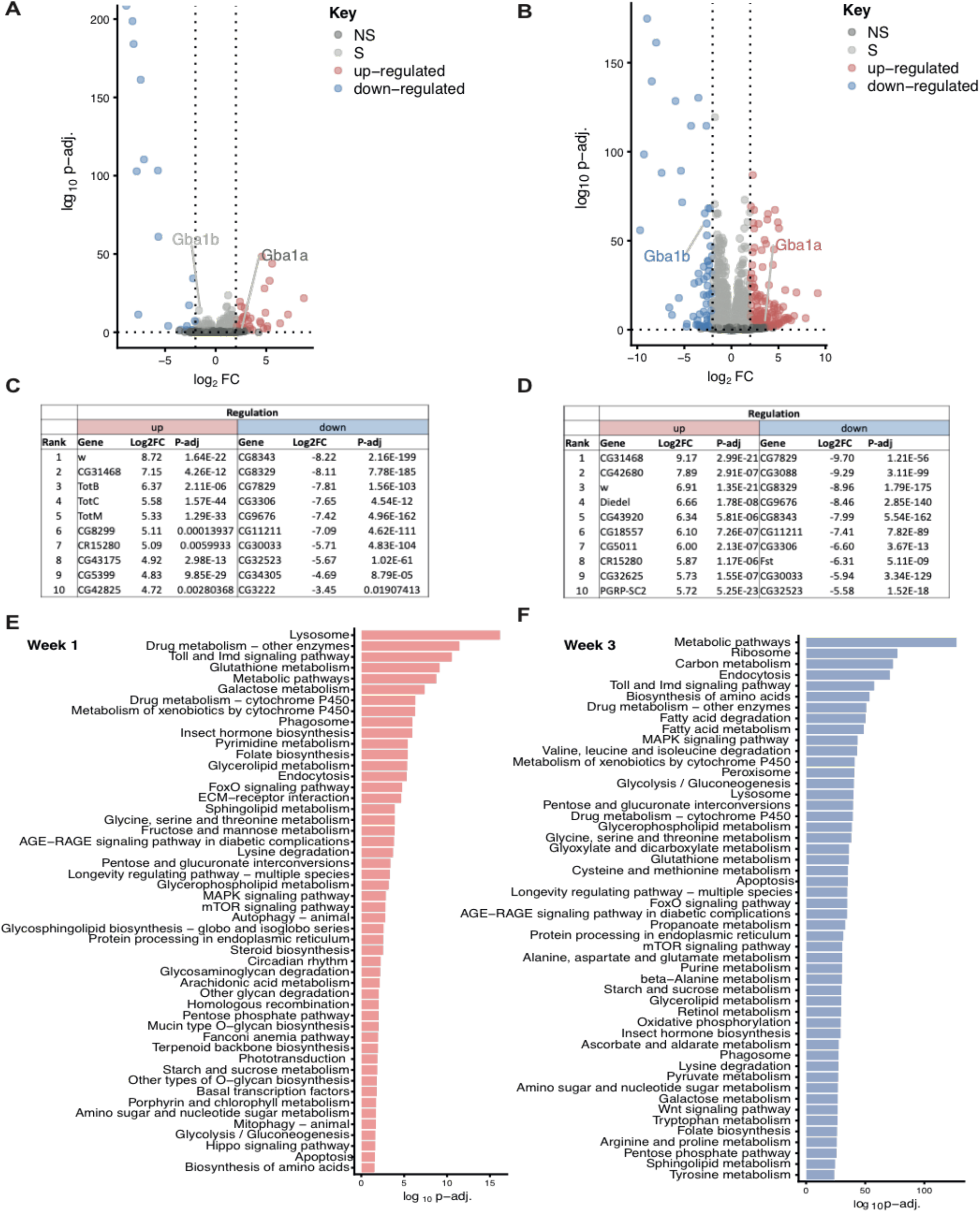
Innate immune system genes are up-regulated in the heads of *Gba1b*^*-/-*^ flies compared to controls. **(A-B)** Volcano plots showing differentially expressed genes in the heads of *Gba1b*^*-/-*^ flies at 1 week **(A)** and 3 weeks **(B)** of age relative to their controls. Differential expression analysis was performed using DESeq2 (https://doi.org/10.1186/s13059-014-0550-8) and identified 11,324 differentially expressed genes at week 1 and 11,883 differentially expressed genes at week 3. Of these genes, 247 and 2545 were significant (S), (p-adjusted <0.05) at 1 week and 3 weeks, respectively. Of the significant genes (light grey) at 1 week, 34 were up-regulated (red) and 18 were down-regulated (blue). The significant genes at 3 weeks comprised of 178 up-regulated and 72 down-regulated genes. The up- and down-regulated genes were selected using the criteria p-adjusted <0.05 and log2 fold change of ± 2 (dotted lines). In addition to the non-significant (NS) genes (dark grey), the two fly *Gba1* genes, *Gba1a* and *Gba1b* are also highlighted and are up-regulated and down-regulated respectively at both time points. **(C-D)** The top 10 up-regulated (up) and down-regulated (down) genes ranked by the log2 fold change (Log2FC) at 1 week (**C**) and 3 weeks (**D**). **(E-F)** The top 50 KEGG pathways of the significant genes at both time points. KEGG pathways of the significant genes at week 1 (247, red) and at week 3 (2545, blue) were obtained using the KEGG database (https://doi.org/10.1093/nar/28.1.27). Top pathways were calculated using the phyper package to compute significance (shown as p-value) between the number of genes in the query and the total genes associated with a particular pathway. Ranking was calculated as -log10 of the adjusted p-value and only the pathways with adjusted p-values <0.05 are shown.

**Figure S2.**
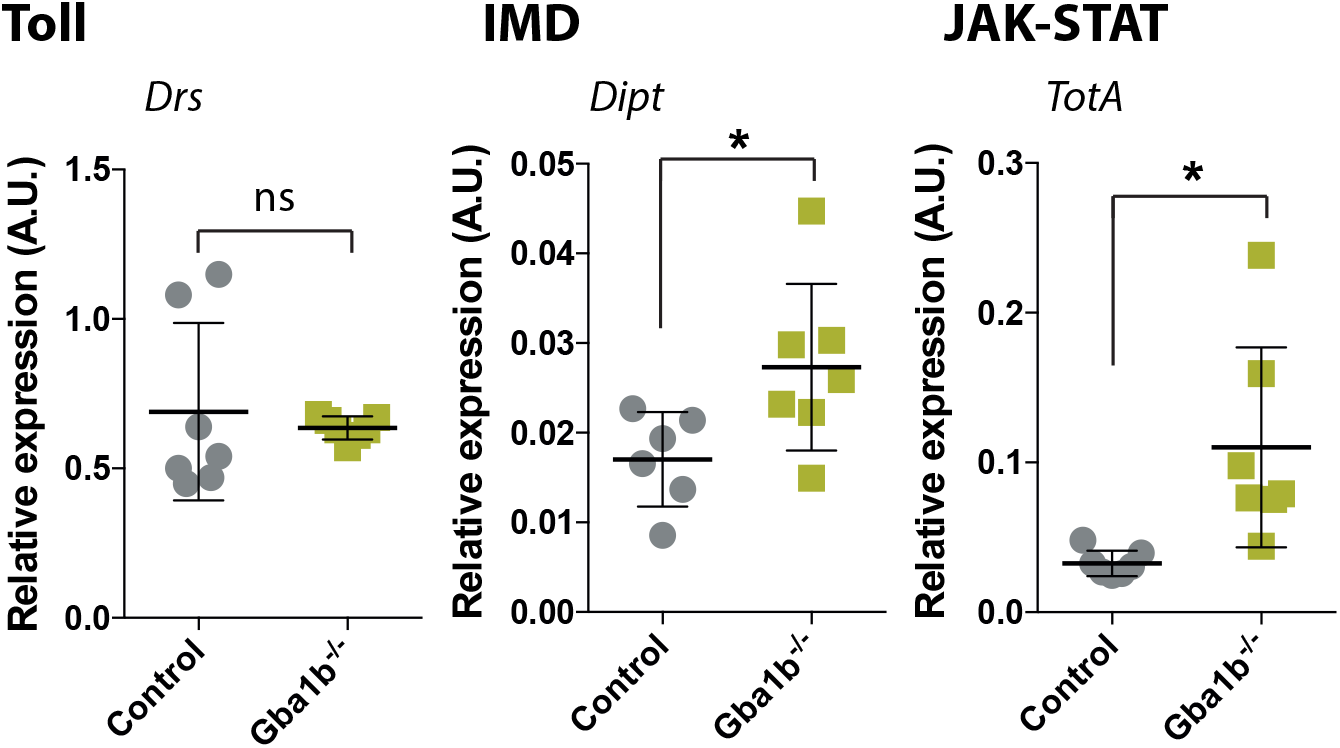
Innate immune pathways are up-regulated peripherally in *Gba1b*^*-/-*^ flies. **(A-B)** Quantitative RT-PCR confirms up-regulation of the IMD (*Dipt)* and JAK-STAT (*TotA*) reporter genes in the headless bodies of 3-week-old *Gba1b*^*-/-*^ flies compared to controls (*Drs*, p=0.64; *Dipt*, *p=0.035; *TotA*, *p=0.0102; n=5-7 per genotype). All target gene expression levels are normalized to tubulin. P-values were calculated using an unpaired t-test and are presented as mean ± SD.

**Figure S3.**
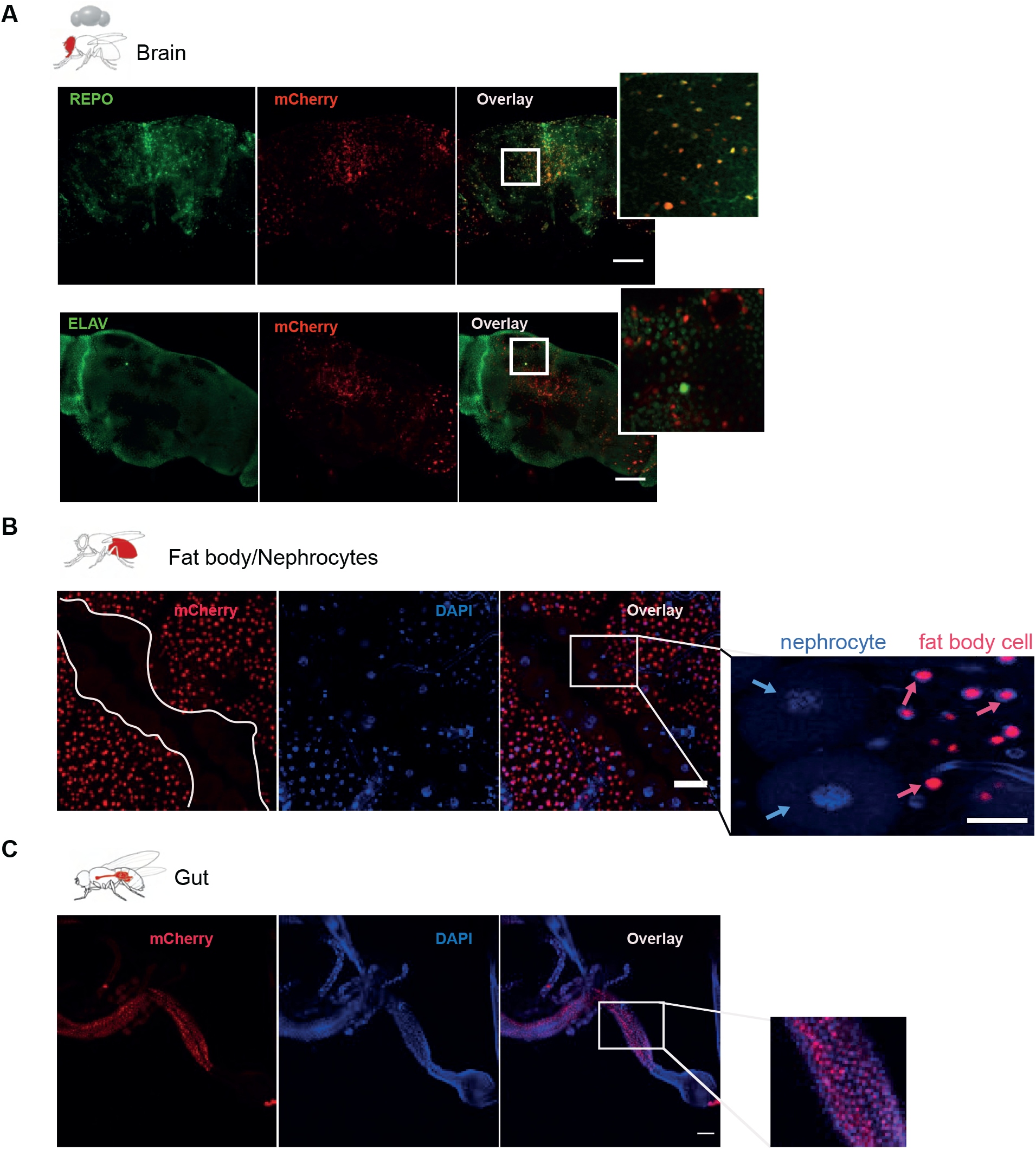
*Gba1b* is expressed in brain, fat body and gut. **(A)** The *Gba1b* gene expression pattern, as assessed by the expression of mCherry under the *Gba1b* endogenous promoter (red channel), overlaps with immunostaining for the glial marker *Repo* (green channel, top panel), but not with the neuronal marker *Elav* (green channel, bottom panel). Scale bars 50 µm. **(B)** Representative confocal image of *Gba1b* expression in abdominal tissues (white lines separate fat body from nephrocyte cells centrally). *Gba1b* expression, as indicated by mCherry expression driven under the *Gba1b* endogenous promoter (red channel), is highly expressed in fat body cells (magenta arrows) but not in nephrocytes (blue arrows). **(C)** *Gba1b* expression in 3-week-old fly gut (red channel). *Gba1b* is highly expressed in the midgut.

**Figure S4.**
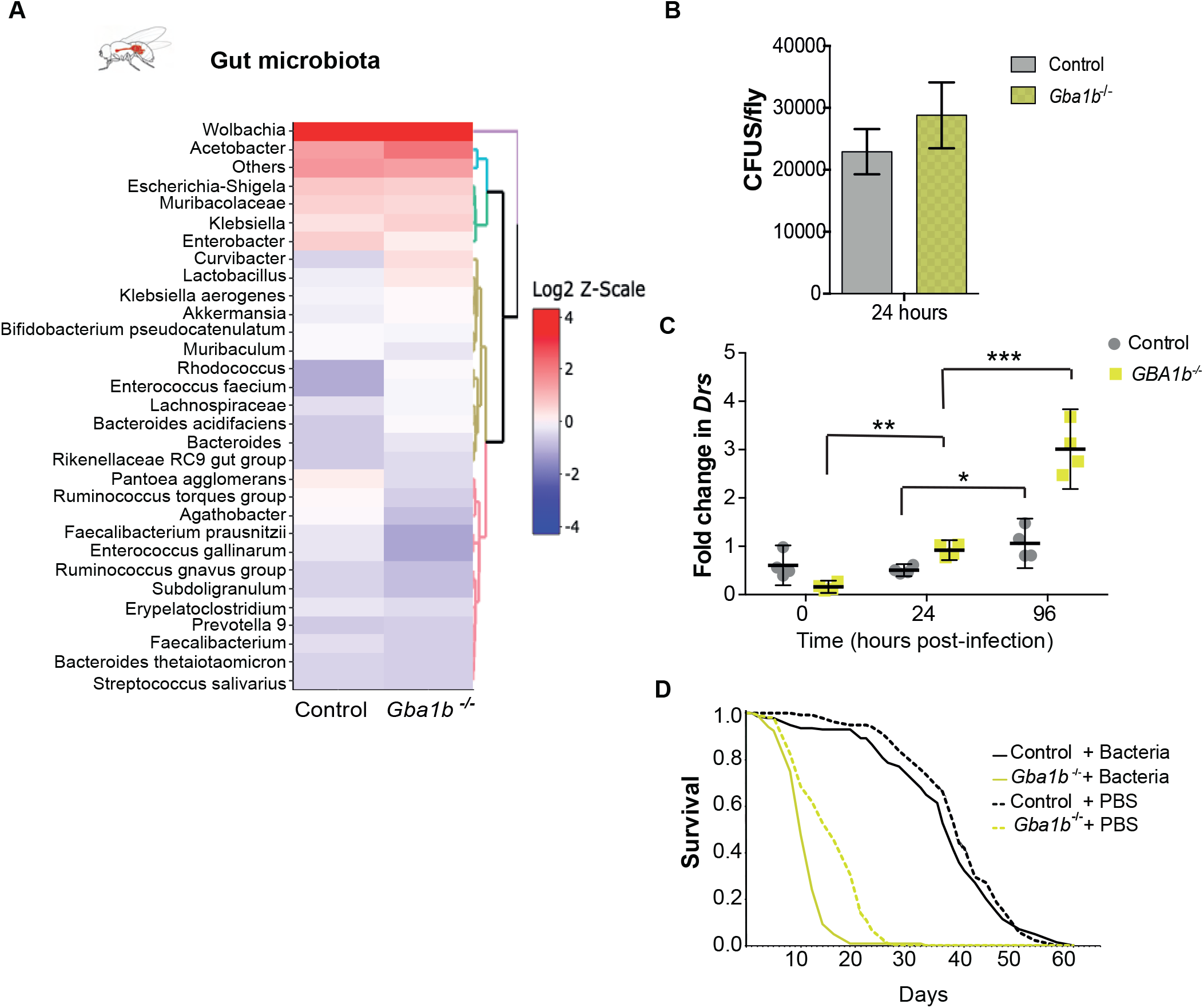
*Gba1b*^-/-^ flies exhibit microbiome dysbiosis. **(A)** Heatmap showing the relative abundance of bacterial genera present in the midguts of *Gba1b*^-/-^ flies compared to their respective controls. Relative abundance is shown as a log2 transformed Z-scale, n=5 per genotype. **(B)** Colony-forming units (CFUs) assay performed on gut tissue to determine the number of *Lactobacillus plantarum* bacteria fed to 3-week-old GF control and *Gba1b*^*-/-*^ flies. Both fly genotypes were fed with similar levels of bacteria (p=0.3702, t-test). **(C)** Whole body *Drs* levels are significantly elevated in *Gba1b*^-/-^ flies compared to controls following an intrathoracic injection of heat-killed *S. aureus* (*Gba1b*^*-/-*^ 0 vs 24 hrs **p=0.00424; 24 vs 96 hrs ***p=<0.0001). Fold change in *Drs* expression between PBS and bacterial injection is shown normalized to tubulin. Two-way ANOVA followed by multiple comparison tests, n=4-6 per condition. Data are presented as mean ± 95% confidence intervals. **(D)** Lifespan of *Gba1b*^*-/-*^ and control flies after an intrathoracic injection of heat killed *S. aureus* or PBS vehicle at 3 weeks of age. The survival curves demonstrate that there is no significant difference in lifespan of control flies treated with either bacterial or PBS injection (p=0.062). *Gba1b*^*-/-*^ flies display reduced survival following heat-killed bacterial injection compared to flies receiving PBS vehicle injection (p=2.5×10^−11^). Log-rank tests were used for all comparisons, n=108-140 flies per condition.

**Figure S5.**
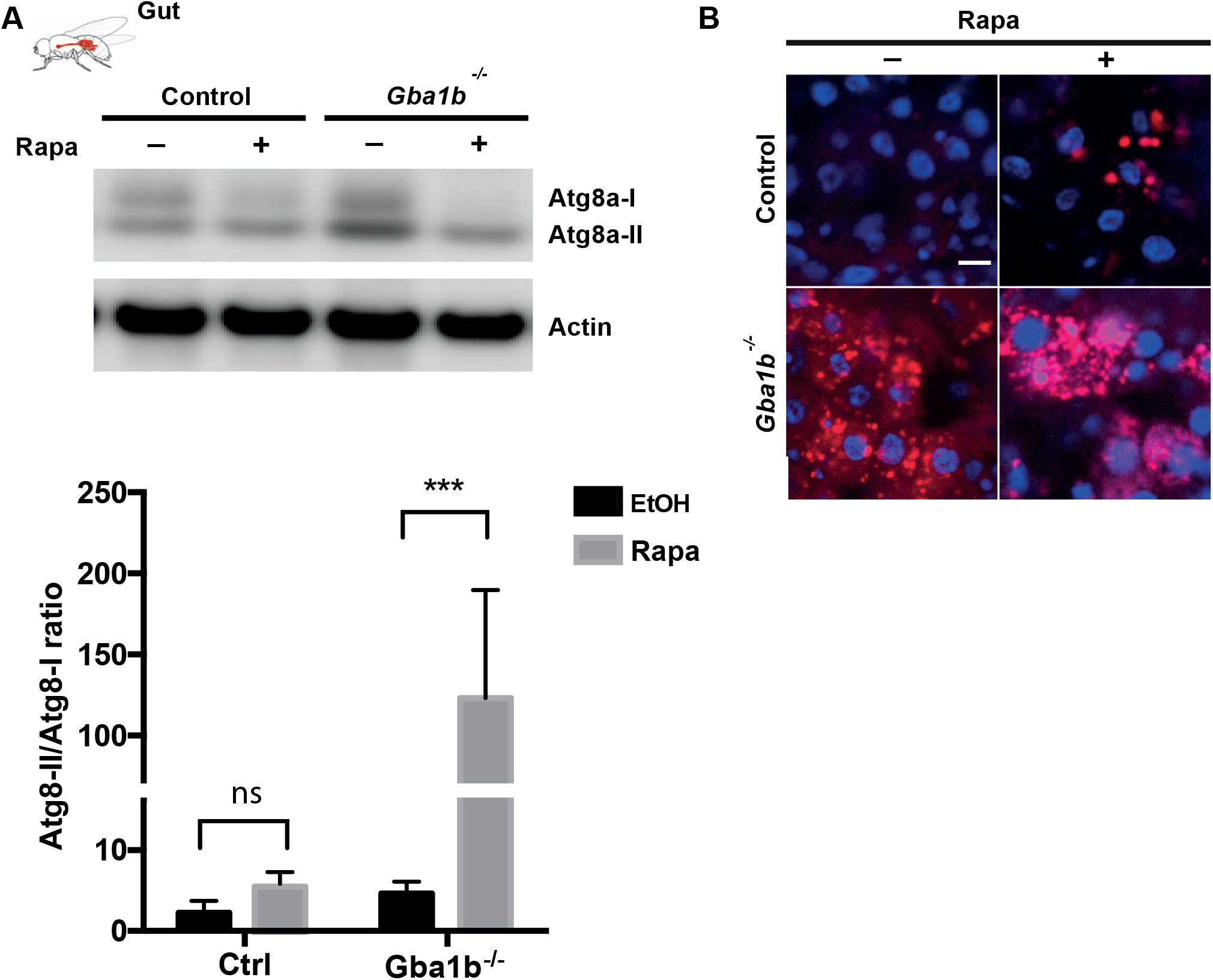
Rapamycin stimulates autophagy in control and *Gba1b*^*-/-*^ flies. **(A)** Immunoblot of autophagy-related proteins, Atg8a-I, Atg8a-II in the fly gut in 3-week-old flies treated with rapamycin (Rapa). Increased ratio of Atg8a-II/Atg8a-I is observed upon drug treatment although this is only significant in the mutant (Two-way ANOVA followed by Fisher’s multiple comparison tests, *Gba1b*^*-/-*^ EtOH vs *Gba1b*^*-/-*^ Rapa ***p=0.001). **(B)** Number of puncta stained by Lysotracker increases in 3-week-old mutant and control flies treated with rapamycin.

## Notes

### Competing Interest Statement

The authors have declared no competing interest.

### Summary of Updates

Author list correction

